# Targeted dynein inhibition generates flagellar beating

**DOI:** 10.1101/153254

**Authors:** Jianfeng Lin, Daniela Nicastro

## Abstract

Motile cilia and flagella are highly conserved organelles that are essential for the normal development and health of many eukaryotes including humans. To reveal the molecular mechanism of motility, we used cryo-electron tomography of active sea urchin sperm flagella to directly visualize the macromolecular complexes and their structural changes during flagellar beating. We resolved distinct conformations of dynein motors and regulators, and showed that many of them are distributed in bend-direction-dependent fashion in active flagella. Our results provide direct evidence for the conformational switching predicted by the “switch-point-hypothesis”. However, they also reveal a fundamentally different mechanism of generating motility by inhibiting dyneins, rather than activating them, causing an asymmetric distribution of force and thus bending. Our high-resolution structural and biochemical analyses provide a new understanding of the distinct roles played by various dyneins and regulators in ciliary motility and suggest a molecular mechanism for robust beating in an all-or-none manner.

**HIGHLIGHTS:**

1. Direct visualization of the switching mechanism for flagellar motility.
2. Flagellar beating is generated by oscillating the side of dynein inhibition.
3. I1 dynein, I1-tether and N-DRC are key regulators that are involved in switching.
4. Structural insights into distinct roles of dynein isoforms in flagellar motility.

## INTRODUCTION

Motile cilia and flagella^*^ are found in nearly all eukaryotes and propel the movement of cells or fluids. Ciliary motility is critical for embryonic development and organ function (Mitchell, 2007), and abnormal motility is implicated in human diseases such as primary ciliary dyskinesia, heart disease, hydrocephalus, and infertility (Afzelius and Stenram, 2006; Fliegauf et al., 2007; Li et al., 2015). Across species, motile cilia share a highly conserved 9+2 axoneme structure with nine doublet microtubules (DMTs) surrounding a central pair complex (CPC) with two singlet microtubules. The DMTs are connected to each other by interdoublet linkers such as the nexin-dynein regulatory complex (N-DRC) (Figure 1A and 1B; Heuser et al., 2009). Ciliary motility is driven by dynein motors, which consist of several polypeptides, including the ~500 kDa heavy chain (HC) with cargo-binding tail, ATP-hydrolyzing motor head and an extended stalk that binds to the microtubule track. In cilia, dyneins are permanently attached to the A-tubule of each DMT through their tail-domains, forming rows of outer and inner dynein arms (ODAs and IDAs) (Figure 1B) (reviewed by Mizuno et al., 2012). ODAs repeat every 24 nm along the length of the DMT and can contain two (e.g. sea urchin, humans) or three (e.g. *Chlamydomonas, Tetrahymena*) dynein HC. IDAs form a heterogeneous group of six single-headed dyneins (IDAs *a-e* and *g*) and one two-headed I1 dynein (IDA *f*) that repeat – like most other axonemal complexes - every 96 nm along the length of each DMT (Figure 1B). Remarkably, each 96-nm axonemal repeat contains more than 400 different proteins (Pazour et al., 2005).

**Figure 1.**
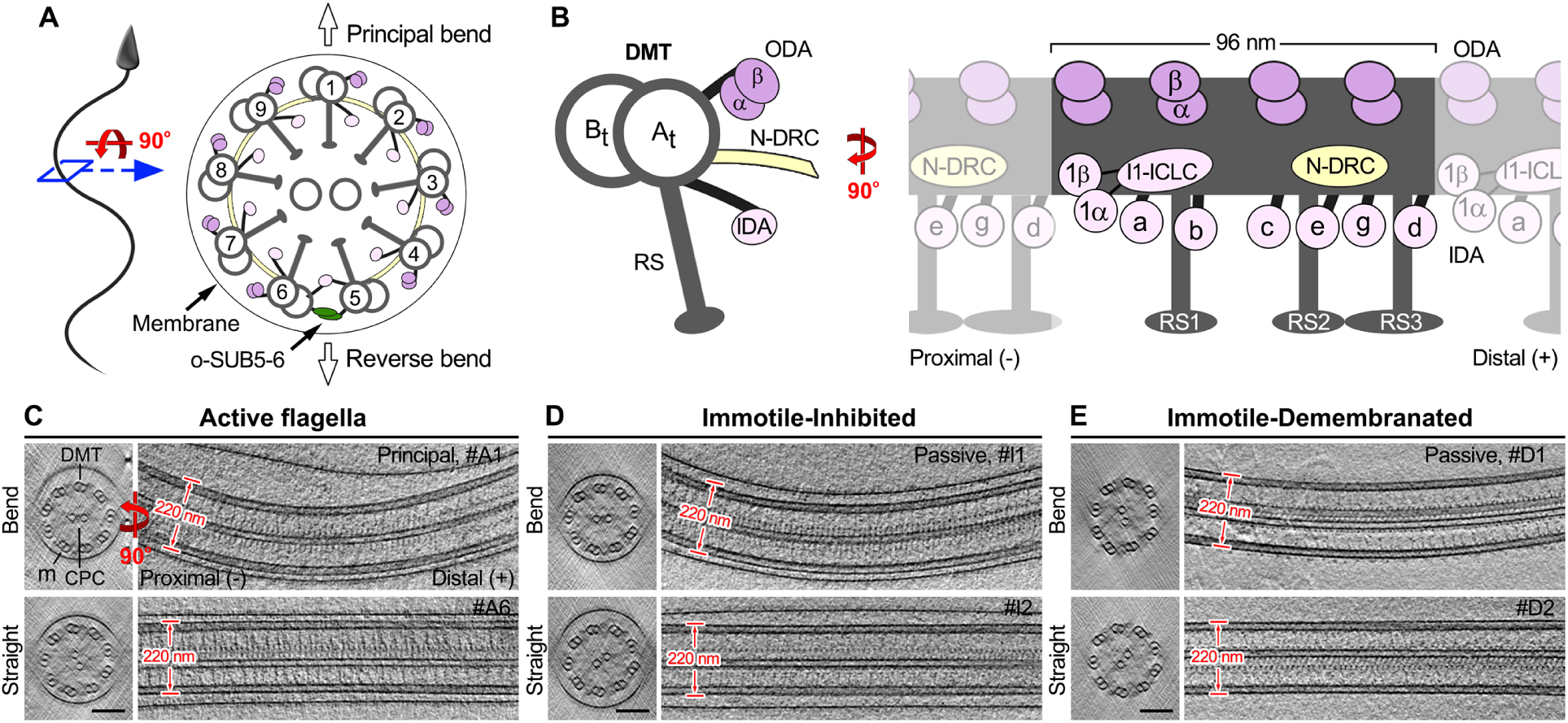
3D visualization of native sea urchin sperm flagella using cryo-ET. (A) Diagrams of a sea urchin sperm and its flagellum viewed in cross-section from the proximal end. The flagellar bending direction is perpendicular to the plane defined by the two central pair complex (CPC) microtubules towards outer doublet microtubule DMT1 (principal bend) or the o-SUB5-6 bridge (reverse bend) (Lin et al., 2012b; Sale, 1986). (B) Cross-sectional and longitudinal views of a DMT. Each DMT is built from a 96-nm unit that repeats along the length of the DMT. Each repeat contains four outer dynein arms (ODAs) with two dyneins (α and β), the inner dynein arm (IDA) I1 with 1α and 1β dyneins and an intermediate-light chain complex (I1-ICLC), and six single-headed IDAs (*a, b, c, d, e*, and *g*). Other labels: A-tubule (A_t_), B-tubule (B_t_), nexin-dynein regulatory complex (N-DRC), radial spoke (RS), microtubule polarity (+ and - end). (C-E) Tomographic slices of representative bent and straight regions from sea urchin sperm flagella in the following states viewed in cross- and longitudinal sections: (C) intact active flagella; (D) intact inhibited immotile flagella; (E) demembranated immotile (ATP-free) axonemes. Scale bar: 100 nm (valid for C-E). See also Movies S1 and S2.

Ciliary motility is generated by dynein motors walking along neighboring DMTs (Sale and Satir, 1977; Summers and Gibbons, 1971). An active dynein on DMT *n* undergoes a mechano-chemical cycle during which ATP-binding and -hydrolysis cause release from the neighboring DMT *n+1*, a head swings towards the MT minus-end, MT rebinding and a powerstroke through which the tail-bound DMT *n* is pulled into the direction of the MT minus-end (Lin et al., 2014). The activity of axonemal dyneins by themselves causes linear sliding between neighboring DMTs, as shown by sliding disintegration experiments of protease-treated axonemes (Sale and Satir, 1977). However in unperturbed axonemes, interdoublet linkers like the N-DRC are thought to restrict this interdoublet sliding, causing the axoneme to bend towards DMT *n+1* (Sale and Satir, 1977; Summers and Gibbons, 1971).

Although several molecular machineries required for ciliary and flagellar motility have been identified, how these components change in response to one another and/or in concert to facilitate movement is poorly understood. Most importantly, the molecular mechanism that causes the beating motion has only been hypothesized, but never been directly demonstrated. Ciliary motility has been modeled for half a century, and the generally accepted concept is that for a flagellum with planar waveform to generate bends in alternating directions, dyneins are thought to be precisely regulated so that only a subset of dyneins are actively walking at any point in time. According to the prevailing “switch-point” hypothesis (Satir and Sale, 1977; Wais-Steider and Satir, 1979), beating is generated by activating dyneins on one side of the axoneme (DMTs 1-4 or DMTs 6-9) (Figure 1A) and then periodically switching dynein activation between opposite sides of the axoneme. This model is supported by indirect evidence, e.g. the extrusion pattern of DMT during ATP-induced sliding disintegration of demembranated cilia (Hayashi and Shingyoji, 2008; Sale, 1986; Tamm and Tamm, 1984). However, to date, the molecular mechanism of ciliary motility has not been visualized directly, and the precise roles of different dynein isoforms and regulators remain largely unknown.

To directly study the conformational changes of axonemal complexes during ciliary beating at high resolution, we rapidly froze swimming sea urchin sperm cells with actively beating flagella, trapping dyneins and other axonemal components in their current conformational states within milliseconds (Lin et al., 2014; Unwin and Fujiyoshi, 2012). The cryo-immobilized, but natively preserved flagella were then imaged by cryo-electron tomography (cryo-ET) and averaged (Nicastro et al., 2006) to determine the 3D structures and conformational states of all major axonemal complexes. Our results reveal that dyneins and several regulators undergo large conformational changes, and that many of these conformations are distributed in a bend-direction-dependent fashion inside the active flagella. Our data directly visualize that flagellar beating is generated by the asymmetry of motor activity between opposite sides of the flagellum, which is caused by regularly switching the sides of selective dynein inhibition, rather than activation.

## RESULTS

Swimming sea urchin sperm were rapidly frozen while their flagella were actively beating (“active flagella”; Movie S1) to trap dyneins and other molecular components in their different conformational states. The native flagella were imaged and 3D-reconstructed (Figure 1C), before extracting, aligning and averaging repeating structures from the tomograms (Nicastro et al., 2006). The subsequent classification of the subtomograms through principal component analysis (Heumann et al., 2011) allowed us to automatically separate different conformations of major axonemal complexes into class averages (Table S1). The conformations were then mapped to their locations in the functional regions of the flagella’s sinusoidal wave to correlate structure with function. As a control, we also analyzed immotile flagella that were inactivated either using the ATPase inhibitor EHNA (immotile-inhibited/immotile-I, Figure 1D; Movie S2; Bouchard et al., 1981), or by demembranating the flagella and washing away the ATP (immotile-demembranated/immotile-D, Figure 1E). Bends observed in the immotile samples were most likely passive bends induced by external force (e.g. liquid flow) during freezing.

### ODA conformations correlate with bend direction in active flagella

Through classification we identified distinct conformations for all axonemal dynein isoforms (Table S2) and the 8-10% of axonemal repeats with the o-SUB5-6 bridge structures (Figure 1A). As previously shown, in some species the ODAs of one DMT are replaced by specialized bridge structures, such as the o-SUB5-6 structure on DMT5 of sea urchin sperm flagella (Lin et al., 2012b; Lindemann and Lesich, 2010). As expected, in immotile controls all α- and β-ODA heads were in the post-powerstroke conformation (Figures 2E-2H, Post), irrespective of whether the inhibited flagella/axonemes were straight or passively bent (Figures 2J, Immotile-I and #I1/2). In contrast, in active flagella the predominant class of ODAs had both motor domains in a pre-powerstroke conformation (Figures 2A-2C, Pre; Movie S3). In prepowerstroke conformations, the dynein heads are moved towards the minus-end of the DMTs relative to the post-powerstroke conformation, and the linker is in the primed (“spring-loaded”) position tilted away from the stalk (compare Figures 2B with 2F; Figure S1). The α-ODA head that is closer to the A-tubule, was in the pre-I conformation with detached MT-binding domain (Figures 2A and 2C; Figure S1A), and the more peripheral β-ODA head in the pre-II conformation with the MT-binding domain re-attached to the DMT (Figures 2A and 2C; Figure S1B) (Lin et al., 2014).

**Figure 2.**
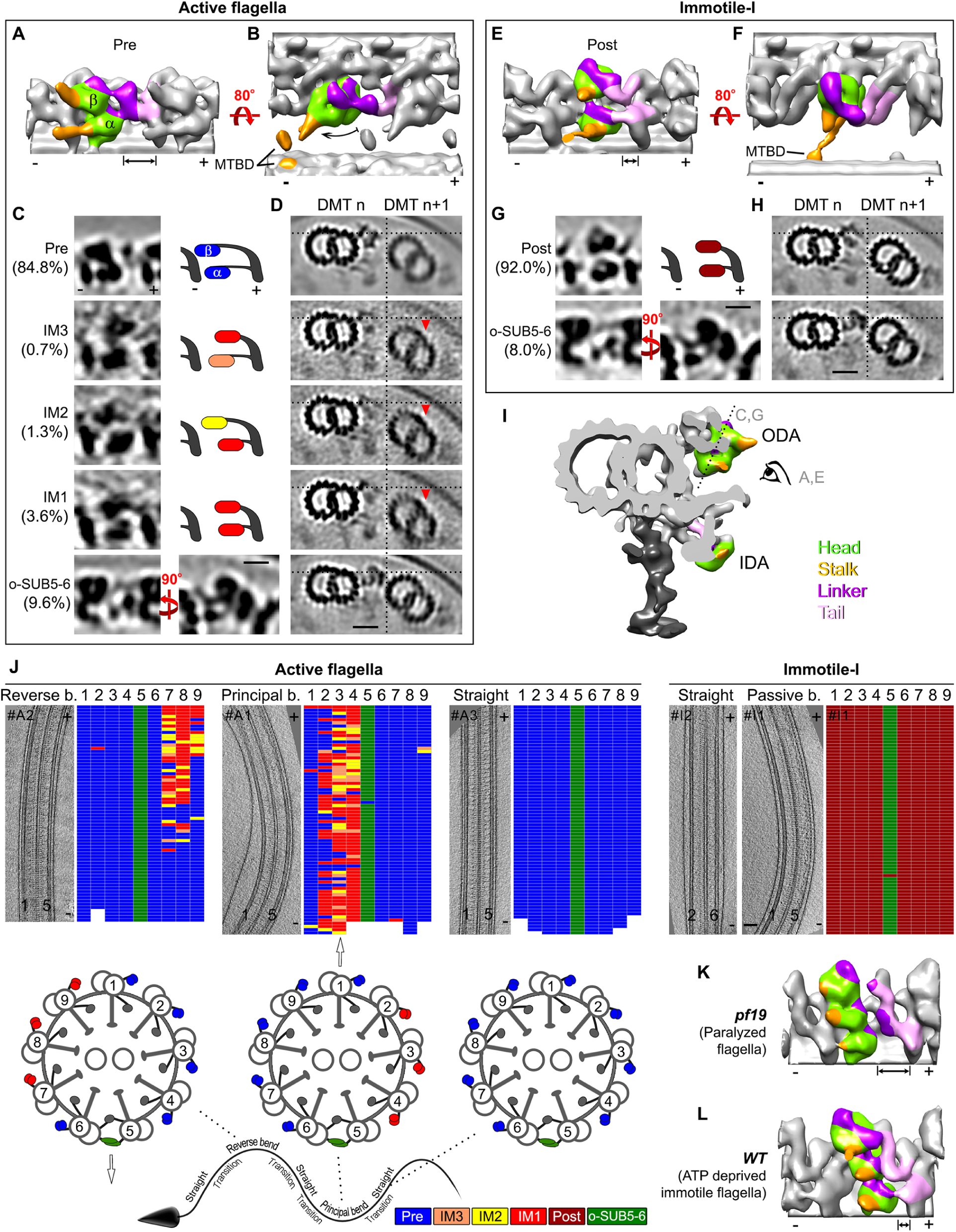
The ODAs exhibit distinct conformations that correlate with bend direction in active flagella (from sea urchin sperm unless otherwise noted). (A and B) 3D isosurface renderings of the class average Pre that has both dynein heads in the pre-powerstroke state. The domains are colored as follows: green (head), orange (stalk with microtubule binding domain, MTBD), magenta (linker), and light pink (tail). The coiled-coil stalks are clearly visible in tomographic slices (see Figure S1, orange arrows). Note that the dynein head is swung towards the microtubule minus-end (-). (C) Longitudinal tomographic slices (left column) of the four averaged ODA classes (Pre, IM3, IM2, IM1) identified in active flagella, and o-SUB5-6 that substitutes the ODA on DMT5. Simplified diagrams (right column) illustrate the changing positions of the two ODA heads in the classes relative to the DMT minus/plus (-/+) ends. The dynein heads are color-coded depending on their conformation as follows: pre-powerstroke (blue, primed and furthest to the minus-end) and intermediate states IM3 (orange), IM2 (yellow) and IM1 (red, which resembles the post-powerstroke state). The percentage of ODA repeats included in each class average is indicated on the left. (D) Cross-sectional tomographic slices of ODA class-averages from active flagella show small positional shifts between neighboring DMTs where ODAs were predominantly in intermediate states (IM1-3). Dotted lines indicate the relative positions of neighboring DMTs, and red arrowheads highlight the shift of DMT *n+1*. (E and F) 3D isosurface renderings of the class average Post that is the major class (92%) in inhibited immotile flagella and has both dynein heads in the post-powerstroke state. Note that both dynein heads are closer to their tails on the microtubule plus side than in the prepowerstroke conformation (compare double-ended arrows in (A) and (E)). (G) Longitudinal tomographic slices (left) of the two averaged ODA classes (Post/dark red, and o-SUB5-6) in inhibited immotile flagella, and simplified diagram (top right) of the two ODA head positions. (H) Cross-sectional tomographic slices of the two ODA class averages in immotile flagella. (I) 3D isosurface rendering of the averaged axonemal repeat of active flagella viewed in cross-section. The location of the tomographic slices shown in (C and G) is indicated by a dotted line, and the viewing direction of (A and E) is indicated by an eye. (J) Distributions of ODA conformations in different regions of individual active and inhibited immotile (Immotile-I) flagella. For each flagellum the following is shown: (top left) a longitudinal tomographic slice (+/- indicates microtubule polarity), (top right) the distribution pattern of ODA conformations, and (bottom, only for active flagella) a diagram of an axonemal cross-section (arrows indicate bend directions). In the distribution patterns, the ODA repeats on the nine DMTs (1-9) are schematically shown as individual grids; the grid color represents the conformational state of each ODA repeat (see bottom legend). (K and L) 3D isosurface renderings of averaged ODAs from paralyzed *Chlamydomonas pf19* mutant flagella (intact cells, K) and isolated WT flagella (with dilution/removal ATP, L). Scale bars: (C and G) 10 nm, (D and H) 20 nm, (J) 100 nm. See also Figures S1-S3 and Movie S3 and Tables S1 and S2.

Intriguingly, the remaining three ODA classes of active flagella showed intermediate conformations (IM1-3), and were restricted to *bent* regions of flagella in a small number of repeats. Importantly, they were found either on DMTs 2-4 in principal bend regions, or on DMTs 7-9 in reverse bends (Figures 2J). In these conformations, at least one of the two dynein heads was in a post-powerstroke-like conformation, and the other head either also in a post-powerstroke-like (Figure 2C, IM1) or in an intermediate conformation where the dynein head was located somewhere between the post- and the pre-powerstroke positions (Figure 2C, IM2/3). Whereas the binding of the dynein stalk to the neighboring DMT was clearly visible in the class averages of pre-II and post-powerstroke dyneins (Figures S1B and S1C), the stalk was not observed in the averages of intermediate or post-powerstroke-like conformations (IM2/3). This suggests that the stalks adopt variable positions in these states, and do not or only weakly interact with the DMTs. In straight regions of active flagella, all ODAs were in the pre-I or pre-II powerstroke conformation (Figure 2J).

The clutch-hypothesis predicts that changes of the axoneme diameter, and thus the inter-doublet distance, are important for ciliary motility, because increased inter-doublet distance could make it difficult for dyneins to attach to the neighboring B-tubule (Lindemann, 2007). Cross-sections of our active flagella show that the axoneme diameter and relative position of DMTs remain mostly the same in both bent and straight flagellar regions (Figure 1C). However, between neighboring DMTs where the ODAs were in intermediate conformations, a small change was observed in the relative position of the DMTs, i.e. here the neighboring doublet (DMT *n+ 1*) was located slightly lower (towards the axoneme center) (Figure 2D). This seems to slightly increase the inter-doublet distance. However, it is not clear if this distance would be prohibitive for dynein binding, as it would depend on which protofilament(s) the dyneins bind to and the angle of the stalk, which was difficult to determine for the intermediate states in the 3D tomograms.

### ODAs are in pre-powerstroke conformations in paralyzed mutant flagella

The observation that the majority of ODAs in active flagella were in pre-powerstroke conformations strongly suggests that axonemal dyneins readily bind ATP, which is omnipresent in the ciliary matrix, and undergo a mechano-chemical cycle by default, i.e. without the requirement to be activated by regulators. So far, dyneins in paralyzed mutant flagella were assumed to be inactive, like in axonemes without ATP. Our data, however, suggests that dyneins in paralyzed mutants with defective regulators, should still exhibit (mostly) primed pre- rather than post-powerstroke conformations as long as ATP is present. To test this hypothesis, we performed cryo-ET of *Chlamydomonas pf19* flagella, which are paralyzed due to the absence of the CPC, a major regulatory complex of ciliary motility (Witman et al., 1978). Indeed, we found that in *pf19* flagella the ODAs were predominantly in pre-powerstroke conformations (Figures 2K, S2C, and S2D) similar to the ODAs in active wild-type (WT) *Chlamydomonas* flagella (Figures S2G and S2H) and active sea urchin sperm flagella (Figures 2A and 2C). This is in contrast to the *Chlamydomonas* controls (i.e. in isolated *Chlamydomonas* flagella and axonemes from which ATP was washed away), where the ODAs were in their post-powerstroke conformation (Figures 2L, S2I-S2P), similar to those of immotile sea urchin sperm flagella and axonemes (Figures 2E-2G).

Our observations present an intriguing model: if flagellar regulation fails, and all dyneins attempted to walk at the same time, the collective pulling forces would be symmetrically distributed across the entire flagellum, counter-balancing each other. The result would be the stiff-immotile phenotype observed for *pf19* flagella, which is interrupted by randomly occurring asymmetry that leads to occasional short-lived twitching, as has been observed for decades (Adams et al., 1981; Hayashibe et al., 1997). Although the ODAs in *pf19* flagella were predominantly in pre-powerstroke conformations, we hypothesize that the tug-of-war between opposite sides of the flagellum could stall many dyneins in their primed states (“catch bond”) and thus prevent futile ATP hydrolysis. This is supported by our comparative measurement of ATP consumption by isolated axonemes from WT and *pf19* flagella. Our results show that the ATP dephosphorylation rate of *pf19* axonemes is less than half (44%) of that of WT axonemes (Figures S2R and S2S). Indeed, dyneins have been shown to exhibit catch bond behavior (Kunwar et al., 2011; Leidel et al., 2012; Mallik et al., 2013; Rai et al., 2013), where up to a threshold increasing load force on the dyneins causes decreasing detachment rates, i.e. the stronger dynein is being pulled in the direction of the cargo, the stronger dynein binds to the microtubule resisting the load.

### IDA *a*, *g*, and *d* conformations correlate with bend direction in active flagella

Our ODA data indicated that intermediate conformation IDAs should also be enriched in bent flagellar regions. To robustly test this, we next performed an auto-classification analysis to detect IDA conformations along the length of the flagella. Consistent with previous studies using DMT-specific averaging (Bui et al., 2009; Lin et al., 2012b; Nicastro, 2009), our classification analysis automatically identified axonemal repeats that lack IDA *a, b, c,* and *e* (on DMT5) and IDA *b* (on DMTs 1 and 9) (Figures 3C-3E, classes “No”; Figure S3), validating the classification method. Consistent with our ODA results, all single-headed IDAs showed predominantly prepowerstroke conformations in active flagella (Figure 3C, Pre; Movie S4), and the post-powerstroke conformation in immotile control axonemes (Figure 3E). Also in accordance with the ODAs, we found that for three of the IDA isoforms, i.e. dyneins *a, g,* and *d*, the distribution of intermediate conformations (IM1/2) was highly correlated with the bent direction in active flagella (Figure 3D). We also observed some small clusters of intermediate conformations in straight regions immediately following significant bends. Although intermediate conformations were also detected for the other IDA isoforms, i.e. dyneins *b, c,* and *e*, these conformations seemed to be more randomly distributed, and a clear switching pattern was not apparent (Figure 3C and D).

**Figure 3.**
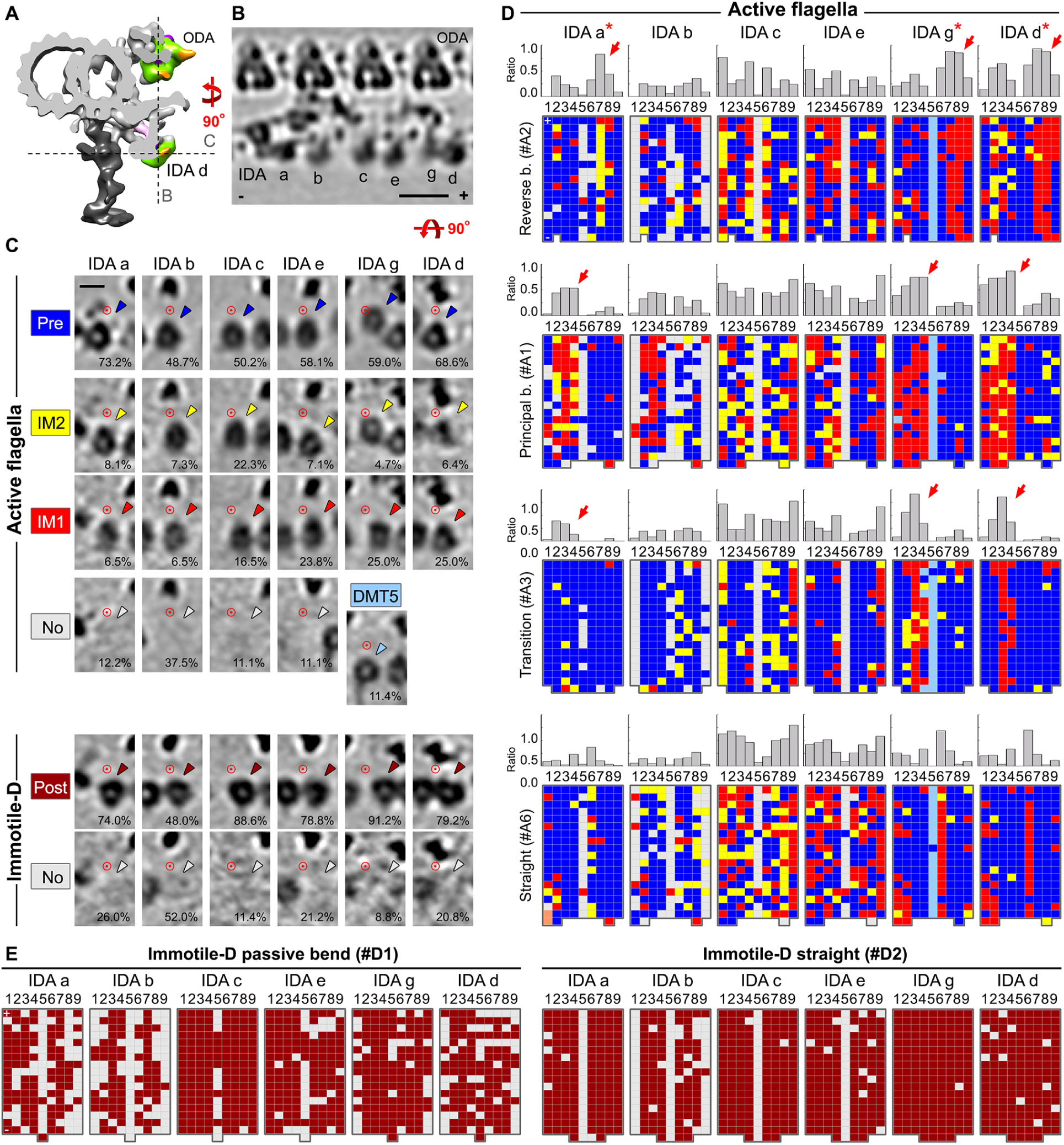
Distinct conformations of single-headed IDAs and their distributions in sea urchin sperm flagella. (A and B) A cross-sectional 3D isosurface rendering (A) and a longitudinal tomographic slice (B) of the averaged 96-nm repeat of active flagella showing the arrangement of IDAs a-e, and g. The dotted lines in (A) indicate the locations of the tomographic slices in (B and C). (C) Tomographic slices of class averages of the single-headed IDAs in active flagella and demembranated, immotile axonemes (Immotile-D). The percentage of sub-tomograms included in each class average is indicated at the bottom of each image. The identified IDA conformations are: pre-powerstroke (Pre, blue), the intermediates IM2 (yellow) and IM1 (red), and post-powerstroke (Post, dark red), a unique conformation of IDA g on DMT5 (DMT5, light blue), and sometimes an IDA was missing (No, gray). Arrowheads point at the particular dynein heads. The red circles mark identical locations in each column to allow better correlation and identification of positional changes of the IDA heads in different conformations; DMT minus end is on the left. (D and E) Distributions of IDA conformations in four functional regions of the flagellar wave of active flagella (D), and in two regions of Immotile-D (E). For each IDA isoform in active flagella (D), an averaged histogram (top) and color-coded distribution pattern of individual flagella (bottom) are shown. The histograms depict the ratios of intermediate states (IM1/2) among all repeats of each DMT. For the reverse bend, principal bend, transition, and straight regions 2, 4, 2, and 5 tomograms were included, respectively. Note that mildly bent flagella were mostly excluded from the histograms due to some ambiguity in assigning them to specific functional regions. The asterisks and red arrows indicate the clustering of IM1 and IM2 conformations in IDAs *a, g,* and *d* in a bend-direction specific manner. For the distribution patterns, the conformations of the IDAs on the nine DMTs (microtubule polarity is indicated with +/-) are schematically shown as individual grids; the grid color represents the conformation of each IDA according to the color-coding in (C). Note that all single-headed IDAs in the Immotile-D are in the post-powerstroke conformation (E). Scale bars: (B) 20 nm, (C) 10 nm. See also Figure S3 and Movie S4.

### Conformations of I1 dynein correlate with bend direction in active flagella

The conformational switching associated with flagellar bends made us next investigate the mechanisms that regulate this phenomenon. The two-headed I1 dynein (dynein *f*) and I1-associated structures are well-known regulators of flagellar motility (Bower et al., 2009; Hendrickson et al., 2004; Toba et al., 2011; VanderWaal et al., 2011). The post-powerstroke conformation of I1 dynein, where both dynein heads and the “intermediate chain and light chain” complex (ICLC) form a planar arrangement of three lobes, was previously described for axonemes (Heuser et al., 2012a), and was also the predominant conformation in the immotile control axonemes here (Figures 4B and 4D; Post).

**Figure 4.**
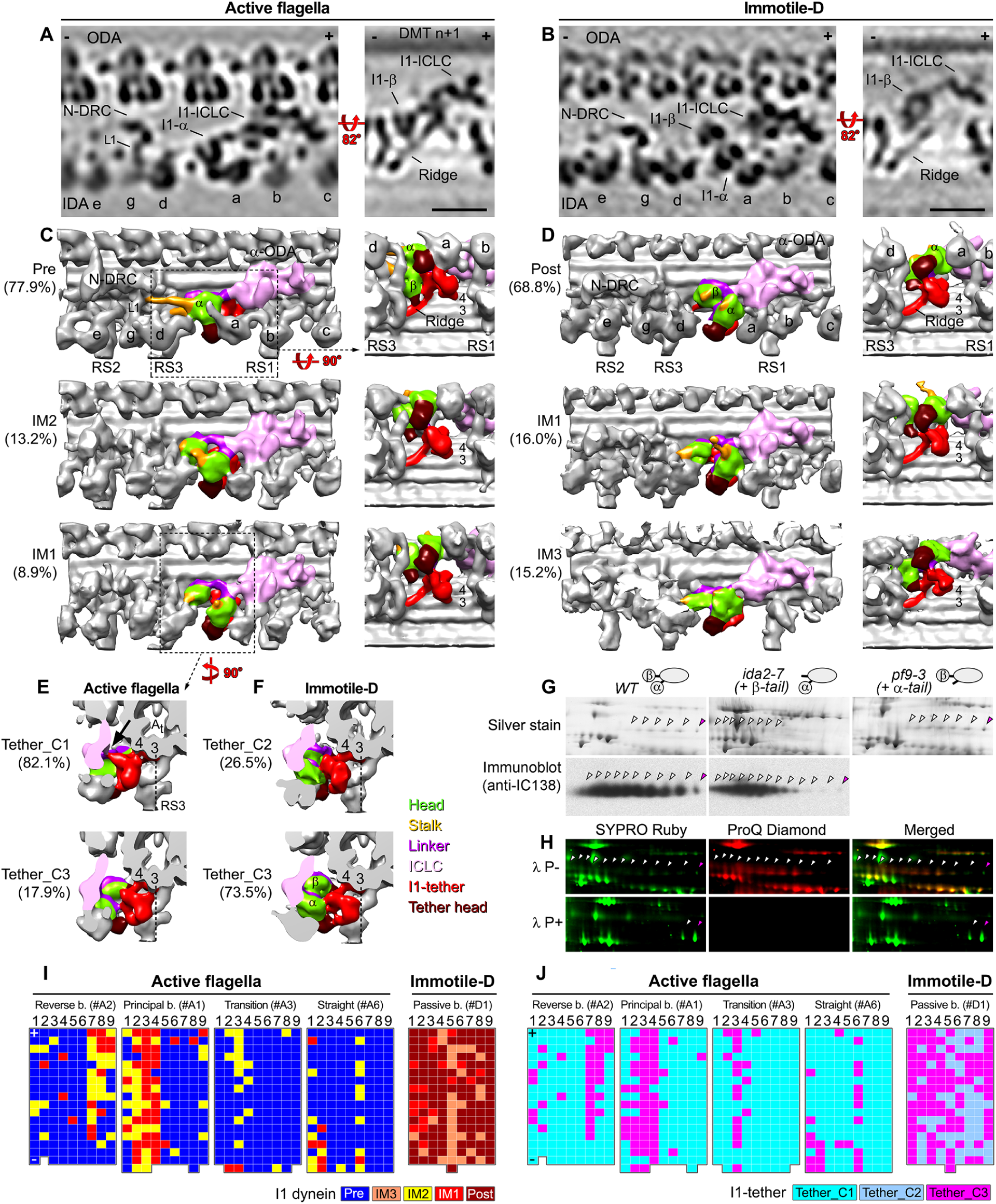
The I1 dynein and I1-tether/tether head complex exhibit several distinct and bend-correlated conformations in active flagella (from sea urchin sperm unless otherwise noted). (A and B) Longitudinal tomographic slices show I1 dynein in its pre-powerstroke conformation in active flagella (A) and in its post-powerstroke state in immotile axonemes (B). Other labels: IDAs a-e, and g; N-DRC, nexin-dynein regulatory complex. (C and D) 3D isosurface renderings of the I1 dynein in the pre-powerstroke (Pre), intermediate (IM1, IM2, and IM3), and post-powerstroke conformations. The following structures are color-coded: I1-tether including its ridge (red), tether head (brown), 1α and 1β dynein heads (green), dynein stalks (orange), linkers (magenta), and the ICLC complex including the dynein tails (light pink). Percentage per class as indicated in the image. (E and F) 3D isosurface renderings (seen in cross-section from distal) show the identified conformations of the I1-tether. Note the interaction between the I1-tether (red) and I1 ICLC complex (light pink) in the Tether_C1 conformation in active flagella (arrow in E). Dotted lines under protofilament A3 serve as reference to highlight the positional changes of the I1-tether in the different conformations. (G) 2DE gels (top) and 2D immunoblots (bottom) of axonemal proteins indicating changes in the phosphorylation of IC138 in *Chlamydomonas ida2-7(+β-tail)* strain that lacks the 1β-motor domain: in WT and *pf9-3(+α-tail)*(a partially rescued I1 mutant that lacks the 1α-motor domain), IC138 exhibits a string of spots (white and magenta arrowheads indicate the phosphorylated and non-phosphorylated isoforms of IC138, respectively); in contrast, in *ida2-7(+β-tail)*, the highly phosphorylated isoforms (on the acidic, left side of the gels) are increased relative to isoforms with a lower level of phosphorylation. (H) 2DE analysis of IC138 phosphorylation in WT *Chlamydomonas* axonemes. The six images show the spot pattern in the absence (*λ P-*, top row) or presence (*λ P+*, bottom row) of λ phosphatase. Phosphorylated isoforms (*white arrowheads*) were detected by both total protein stain (SYPRO Ruby) and phosphoprotein stain (ProQ Diamond). Non-phosphorylated isoforms (*magenta arrowheads*) were only detected by total protein stain. Merged images facilitate the localization and quantification of phosphorylated protein isoforms. (I and J) Distributions of conformations of I1 dynein (I) and I1-tether (J) in different regions of active flagella and immotile axonemes. The I1 dynein and I1-tether on the nine DMTs are schematically shown as individual grids; the grid color represents the conformations according to the color-legend at the bottom. Scale bars: (A and B) 20 nm. See also Figure S3 and Movie S5.

In active flagella, the predominant I1 dynein conformation had both the 1α- and 1β-HCs in pre-powerstroke positions, and exhibited numerous major structural rearrangements (Figures 4A and 4C; Pre) compared to the post-powerstroke state. The dynein heads were shifted away from the ICLC towards the minus end of the DMT – as expected for the Pre-state, but in addition, the two heads underwent a rotation relative to each so that the 1α-head was closer to the neighboring DMT and the 1β-head closer to the A-tubule “hidden” behind the 1α-head, and the stalks pointed proximal towards the N-DRC, rather than to the neighboring DMT. In fact, the MT-binding domain of the 1α-head appeared connected to the base of the N-DRC L1-linker arm (Figure 4C; Pre).

Interestingly, the intermediate states IM1 and IM2 (Figure 4C; IM1 & IM2), showed a bent-direction specific distribution in active flagella. Similar to the ODAs and IDAs *a, g,* and *d*, the I1 dynein IM1 and IM2 conformations were primarily clustered on DMTs 2-4 in the principal bend, and DMTs 6-9 in the reverse bend (Figure 4I). In addition, the region with intermediate states extended often into the straight regions just passed significant bends (Figure 4I, transition).

### Major conformational changes of the I1-tether show switching pattern in active flagella

We previously identified an I1 dynein associated structure that surprisingly tethers the I1 motor domain to the “cargo” A-tubule (Heuser et al., 2012a; see also Figures 4A-4E). We named the two domains of this structure the “tether head” that is attached to the dynein head, and the “tether” that links the tether head to the A-tubule (Heuser et al., 2012a). Here, we show that this complex undergoes bend-correlated major structural changes in active flagella that suggest an important regulatory function.

In immotile controls, two distinct I1-tether conformations (tether_C2 and the predominant tether_C3) were observed (Figure 4F), but without obvious correlation with bend direction (Figures 4J and S3B). In contrast, the two conformations found in active flagella, tether_C1 and tether_C3, showed a bend-correlated distribution. The tether_C3 conformation (Figure 4E) was highly correlated with the intermediate conformations IM1 and IM2 of I1 dynein (Figure 4C), i.e. tether_C3 primarily clustered on DMTs 2-4 in the principal bend, and DMTs 6-9 in the reverse bend (Figure 4J), including extending into the straight regions just passed major bends (Figure 4J, transition). The predominant tether_C1 conformation, in which the I1-tether was rotated away from protofilament A3 and connected to the ICLC complex (Figure 4E; arrow), was found almost exclusively in conjunction with the pre-powerstroke I1 dynein (Figure 4C; Pre). Intriguingly, in the tether_C1 state the 1β-tether head connected to the here newly resolved I1-tether ridge on the A-tubule (Figures 4C; Pre; Movie S5) that appears to contact to the base of radial spoke RS3. Similar to the CPC, radial spokes are thought to be upstream regulators that are important for ciliary motility, because radial-spoke defects often result in paralyzed flagella (Brokaw et al., 1982; Warner and Satir, 1974). Although we did not observe radial spoke changes that correlated with specific functional regions of the flagellar wave (Figure S4), the state-specific attachment and dissociation of the tether head from the ridge, implies that RS3 regulates and/or receives signaling feedback from I1 dynein. Interestingly, the RS3 base is known to contain subunits of the Calmodulin- and spoke-associated complex (CSC) (Urbanska et al., 2015); which itself is a regulatory hub that connects between RS2, RS3 and the N-DRC (Heuser et al., 2012b).

Previous studies of *Chlamydomonas* flagella have shown that the intermediate chain IC138 in the I1 ICLC (Heuser et al., 2012a) is an important phosphorylation switch that regulates dynein activity and flagellar motility. In the phosphorylated state of IC138, I1 is inactive, whereas IC138 dephosphorylation restores activity (Bower et al., 2009; Hendrickson et al., 2004; Toba et al., 2011; VanderWaal et al., 2011). Based on our observation of bend-correlated conformational changes in I1 dynein and I1-tether, we hypothesized that the tether-ICLC connection and the interaction between 1β-head and I1-tether ridge play important roles in regulating I1 activity through the IC138 phospho-switch. To test this hypothesis we conducted phosphorylation analysis of two I1 mutants that lack the motor domain of either 1α-HC (*pf9-3*+α-tail) or 1β-HC (*ida2-7*+β-tail) (Myster et al., 1999; Perrone et al., 2000). Indeed, we found that the absence of the 1β-head, but not of the 1α-head, resulted in hyper-phosphorylation and thus inactivation of IC138 (Figures 4G and 4H), consistent with a role for I1 tether and 1β-HC in I1 phospho-regulation.

### N-DRC shows bend-correlated conformational changes, but no significant stretching in active flagella

We previously demonstrated that the nexin link connecting neighboring DMTs is part of the dynein regulatory complex (DRC), now called N-DRC (Heuser et al., 2009). In contrast to a previous hypothesis that the nexin link might stretch to accommodate inter-doublet sliding during beating (Warner, 1976; Olson and Linck, 1977), we could not find any stretching or tilting of the N-DRC in active flagella. However, we did observe small positional and conformational changes of the N-DRC that correlated with the bent direction (Figure 5). For the immotile controls, the classification analysis showed only two distinct classes for the nexin link, the predominant N-DRC_2 that adopted a position in the middle between the rows of outer and inner dyneins as described previously (Figure 5; see also Heuser et al., 2009). The second class, N-DRC_4, was solely associated with DMT5 that contains doublet-specific features such as the o-SUB5-6 bridges and lack of regular IDAs c and e (Figure 5; see also Bui et al., 2009; Lin et al., 2012b). A unique feature of N-DRC_4 was an additional density that extended from the N-DRC distal lobe to the neighboring DMT (Figure 5B).

**Figure 5.**
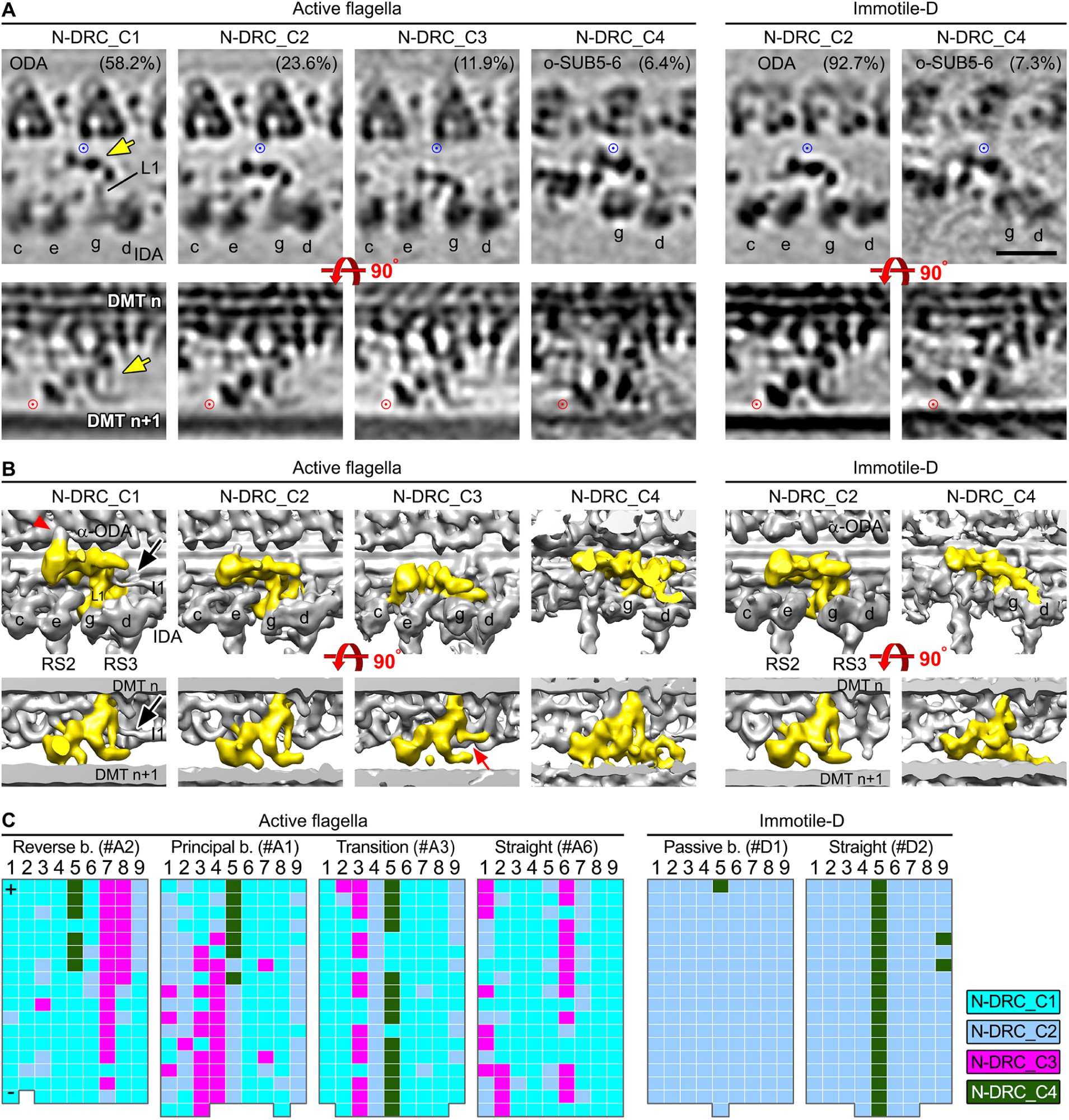
The N-DRC shows bend-correlated conformational changes, but no significant stretching in active sea urchin sperm flagella. (A) Longitudinal tomographic slices of the nexin-dynein regulatory complex (N-DRC) class averages (proximal is on the left). The blue and red circles indicate the same location within each row of images and serve as reference points to facilitate comparison of the N-DRC conformations. Note that none of the N-DRC conformations shows any stretch or tilting in the proximal/distal direction (see position relative to the red circle). Other labels: L1, linker arm L1 that connects from the nexin linker to the motor domain of dynein g. (B) 3D isosurface renderings of averaged N-DRC conformations in the same orientation as in (A). In the N-DRC_C1 conformation, the nexin linker was in close proximity of the ODA row (also see distance from blue circle in A) and forms two unique connections with dyneins: the proximal lobe and distal region of the nexin linker interacted with the stalks of α-ODA (red arrowhead) and I1α-dynein (black arrow), respectively. In N-DRC_C3, both connections were lost, and the distal region rotated ~ 90° and attached to the tail of IDA *g* (red arrow). N-DRC_4 has additional densities extended from the distal lobe. Although it differed slightly between active and immotile flagella, no bend-specific correlation was identified for these differences (C). (C) Distributions of N-DRC conformations in different regions of active flagella and demembranated immotile axonemes. The N-DRCs on the nine DMTs are schematically shown as individual grids; the grid color represents the different conformational classes (as indicated in the legend on the right). Scale bar: (A) 20 nm. See also Figure S3.

In active flagella, the nexin linker adopted a range of positions, but overall they could be clustered into four classes, the previous two plus two additional classes, the predominant N-DRC_1 and N-DRC_3 that were unique to beating flagella. The N-DRC_1 class correlated well with pre-powerstroke dyneins and was shifted closer to the ODAs than the other classes. In addition to the usual L1-linker connection to the motor domain of dynein g, N-DRC_1 showed two more direct connections to dyneins: The MT-binding domain of α-ODA connected to the proximal lobe of the N-DRC (Figure 5B, red arrowhead) and the MT-binding domain of the prepowerstroke I1α dynein was connected to the L1-linker of the N-DRC (Figure 5B, black arrows). In contrast, the N-DRC_C3 class has a low abundance, its nexin linker was shifted closer to the row of IDAs, and the proximal and distal lobes formed direct connections with IDAs *c, e* and *g* (Figure 5B). Importantly, the N-DRC_C3 class was the only state that was clearly distributed in a bend-direction-dependent fashion, i.e. primarily clustered on DMTs 2-4 in principal bend, DMTs 7-9 of the reverse bend, as well as in some straight regions bordering major bends (Figure 5C). Thus the distribution is similar to those of intermediate states IM1 & IM2 of IDAs *a, g, d*, and I1 dynein, and of I1-tether_C3.

## DISCUSSION

Cellular processes are dynamic and complex, requiring many molecular components to self-organize and work together to produce a cellular behavior. Here we demonstrate that comparative cellular cryo-ET studies provide the conceptual framework and experimental tools to better understand cellular function. Our approach revealed the spatial and temporal information required to unravel both the functions of individual building blocks, as well as their interactions and molecular mechanisms that give rise to the emergent mechanics and complex behaviors of larger-scale biological structures, such as cilia. The study revealed unprecedented views of native dyneins and their regulators in different functional states *in situ*. The identification of biologically meaningful spatiotemporal patterns of discrete conformational states of motors and regulators in different functional regions of beating flagella revealed a switch-inhibition mechanism for flagellar motility. This molecular model explains not only the mechanics of how cilia and flagella generate motion, but also how cells can robustly maintain a highly dynamic yet long-lasting process that requires coordinated action of thousands of macromolecular machines and conformational switching in a hundreds of a second.

### The switch-inhibition model for the molecular mechanism of ciliary motility

Our present study provides the first direct visualization of the conformational switching that occurs along opposing sides of a beating flagellum. Unlike the original switch-point hypothesis that predicted the selective activation of dyneins on opposing flagellar sides, we observed that most dyneins exist in “spring-loaded” pre-powerstroke conformations, whereas only a few motors were in inactive intermediate and/or post-powerstroke-like states that were not or only weakly bound to the neighboring DMT (Figures 2-4). The prevalence of active-state dyneins is consistent with the fact that most isolated dyneins can readily undergo a mechanochemical cycle *in vitro* in the presence of ATP+Mg^2+^ without requiring additional activating proteins (Imamula et al., 2007; Kagami and Kamiya, 1992; Kon et al., 2005; Tsygankov et al., 2009). It is also supported by the fact that most paralyzed flagellar mutants (e.g., *pf19*) can undergo ATP-induced, dynein-driven sliding disintegration after proteolytic cleavage of inter-doublet linkers (Witman et al., 1978). Therefore, our observations strongly imply that dyneins are active by default, and ready to undergo the mechanochemical cycle as long as ATP is present. In turn, *selective inhibition* (rather than activation) generates the required asymmetry of motor activity and force between opposite sides of the flagellum, resulting in flagellar beating. We name this new model for flagellar motility the switch-inhibition model.

Given that bends are initiated at the flagellar base and travel over time to the tip, the sequential conformational changes we observed along frozen flagella also represent the temporal sequence of structural changes during wave formation. By correlating structural changes with their location in the flagellar bend, we are able to propose the following switch-inhibition model for the generation of ciliary and flagellar motility (Figure 6, Movie S6):

**Figure 6.**
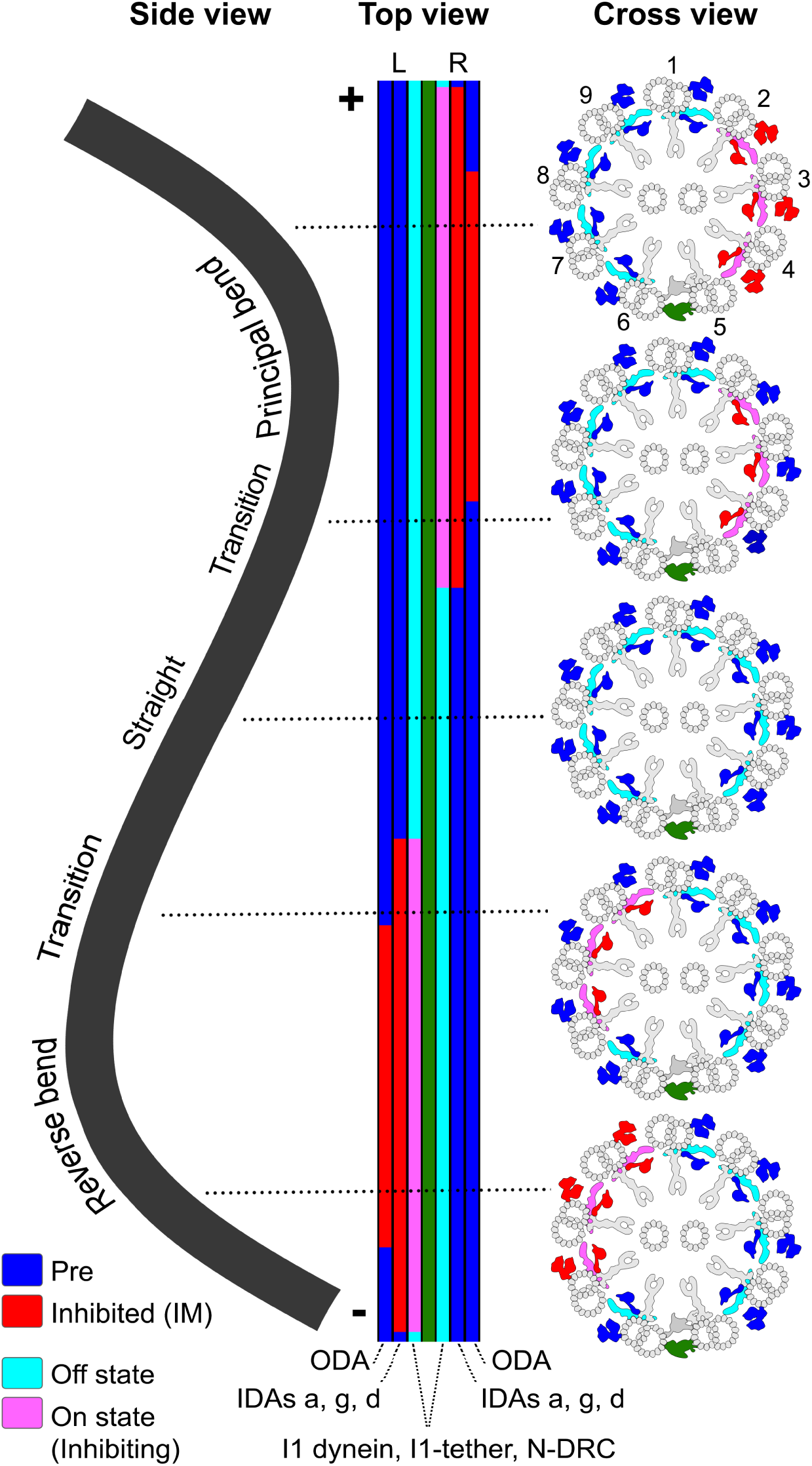
Schematic model of the switch-inhibition mechanism of ciliary/flagellar motility. Summary illustration of a sinusoidal wave of an active flagellum that contains reverse bend, transition, straight, and principal bend regions. For explanation of the switch-inhibition mechanism please see the Discussion. The schematic shows the flagellum in different views: (left) longitudinally from its “right” side (where DMTs 2-4 are located), (middle) viewed from the top passing through DMT1 and o-SUB5-6 bridge (L/R indicate the “left” and “right” sides of the flagellum), and (right) cross-sectional views from proximal. In the top and cross-sectional views, the distributions of the different conformations of various axonemal complexes (as indicated at the bottom of the top view) are shown in the different functional regions of the flagellum. The different conformations are indicated by distinct colors (as specified in the color-legend at the bottom left). See also Figure S3 and Movies S3-6.

**a) Straight regions:** Under physiological conditions (ATP present) and with the regulators in their “off” state or the inhibitory signal still below a threshold, the dyneins on DMT *n* undergo mechanochemical cycles and generate force through transient, ATP-dependent interactions with the B-tubule of DMT *n+ 1*. However, because the nine DMTs are arranged radially symmetric in a cylinder, the dynein forces from opposite sides of the axoneme are counterbalanced, keeping the axoneme straight. The opposing forces exert a resisting load on the dyneins, causing them to stall, halting their mechanochemical cycle (incl. ATP-hydrolysis), which may lead to stronger microtubule-binding through catch bond formation (Figure 6, the 3rd cross section from top to bottom). Recent studies of cytoplasmic dyneins have indeed reported that dyneins are catch-bond motors (Leidel et al., 2012; Rai et al., 2013) similar to myosin motors (Guo and Guilford, 2006; Thomas et al., 2008). Without proper function of regulators that can “break” the force balance, flagella appear paralyzed and stiff, such as the here studied *pf19* mutant flagella.

**b) Bend-initiation:** When signals switch regulators (such as the N_DRC, I1 dynein and I1 tether/tether head) to their “on” (inhibitory) states, they specifically inhibit the activity of IDAs *a, g,* and *d* on either DMT2-4 or DMT7-9. As long as the effect is below the threshold for “catastrophic” release of all dyneins on the inhibited side, the net force generated by dyneins on the opposite side is not yet sufficient to overcome the stiffness of the axoneme and generate bending (Figure 6, 4^th^ cross sections). When the inhibitory signals reach a supra-threshold level, enough IDAs on one side are inhibited, allowing the dyneins on the opposite side to be both released from catch bond and to exert a net force sufficiently strong to overcome the stiffness of the axoneme and initiate a mild bend in the flagellum.

**d) Bending:** The positional changes of DMTs caused by the mild bend initiation and/or transmission of the inhibitory signal downstream to the ODAs on the inhibited side of the axoneme (DMT2-4 or DMT7-9) allow for a rapid increase of net force generated by the actively walking dyneins on the opposite side of the flagellum. This causes further sliding between neighboring DMTs, which is restricted by molecular inter-doublet linkers and thus causes a full bend of the flagellum in one direction, i.e. the principal or the reverse bend (Figure 6, 1^st^ and 5^th^ cross sections).

After the inhibitory signals are turned off by so far unknown trigger(s), dynein activity on the previously inhibited side recovers, causing the flagellum to straighten (Figure 6, 2^nd^ cross sections) and the bending cycle restarts. By periodically switching the side of dynein inhibition, the flagellum alternates the direction of bending, resulting in the typical (quasi-)planar waveform of sea urchin sperm flagella.

### Switch-inhibition as robust mechanism for maintaining waveform amplitude

The switch-inhibition model provides a comprehensive molecular mechanism for flagellar beat generation that appears more robust than a switch-activation model. Sea urchin sperm flagella beat with fairly consistent wave amplitude and beat frequency of ~50 Hz (Eshel et al., 1990), i.e. the bending direction switches every ~10 msec. For a “switch activation model”, the regulators would have to activate dyneins in a temporarily coordinated manner with high efficiency, and the bend amplitude would directly depend on the number of synchronously activated dyneins. However, a switch-inhibition mechanism could happen in an “all-or-none” fashion: meaning the combination of targeted inhibition of some dyneins on one side of the flagellum, the resulting load-release from catch bond of the dyneins on the opposite side and a slight increase of the inter-doublet space on the inhibited side (caused possibly by N-DRC conformational change and/or mild bending) could cause cooperative or “catastrophic” microtubule release of the dyneins on the inhibited side, allowing quasi 100% of the inherently active dyneins on the opposite side to rapidly drive bending. Following the all-or-none law, as long as the inhibitory signal and percent of inhibited dyneins exceed a specific threshold, the axoneme would bend consistently with 100% wave amplitude, resulting in the relatively constant and sustained beating of cilia and flagella. “All-or-none” mechanisms are often observed in cellular processes that require rapid switching and/or preservation of signal strength, such as transduction of action potentials and heart muscle contraction (Pareti, 2007; Wacke and Thiel, 2001; Weiss et al., 1976).

### Distinct roles for different axonemal dyneins

It is intriguing that typically a single cytoplasmic dynein isoform is sufficient to fulfill different roles in a plethora of cellular processes from retrograde transport to mitosis, whereas cilia require more than a dozen different axonemal dynein isoforms for proper motility (Hook and Vallee, 2006).. We observed temporal and conformational heterogeneity and specializations among the dynein isoforms, suggesting that they have distinct functions during motility generation. The intermediate conformations of IDAs *a, g, d*, and I1 dynein in transition regions (Figure 3D and 4I) seem to precede the ODA conformational switching (Figure 2J), suggesting that they may play a role in initiating flagellar bending. In turn ODAs, likely add power and speed to the all-or-none reaction, which could influence the beat frequency (Figure 6). This interpretation is consistent with studies of *Chlamydomonas* IDA and ODA mutants, in which the flagella that lack IDA *c* have wild-type beat frequency and only slightly slower swimming velocity (Yagi et al., 2005), whereas flagella that lack IDAs *a, c, d,* and I1 dynein are paralyzed, but can both beat after externally inducing a bend and undergo DMT sliding *in vitro* at near wild-type velocities (Hayashibe et al., 1997; Kurimoto and Kamiya, 1991). In contrast, ODA mutant flagella are motile with nearly normal waveforms but exhibit reduced beat frequency and DMT sliding velocities *in vitro* (Mitchell and Rosenbaum, 1985; Porter et al., 1992; Seetharam and Satir, 2005). The conformational distributions of IDAs *b, c,* and *e* lacked obvious correlation with the bending direction (Figure 3), making these dynein isoforms likely not essential for bend initiation. However, they could still play roles in regulating flagellar waveform, a function that previous mutant studies have attributed to IDAs in general (Kamiya et al., 1991; Perrone et al., 2000; Yagi et al., 2005). A recent study about possible feedback mechanisms for generating the periodic bending motion typical for cilia and flagella, proposed that the dynamic and static mode of the waveform could be independently controlled (Sartori et al., 2016). Therefore, some of the IDA isoforms may e.g. control changing between asymmetric waveform (regular forward swimming) and symmetric waveform (backwards swimming due to photoshock or *mbo2* mutant) of *Chlamydomonas* flagella by changing the static mode.

### Models for periodic conformational switching and outlook

Here we present direct evidence for a switch-inhibition model for ciliary beating and show that dyneins and regulators periodically switch conformations. However, the nature of the switching signal(s) is still unknown. Several models for what causes switching have been suggested, such as dyneins being inherent oscillating force generators (Shingyoji et al., 1998), the distributor model that proposes a regulatory enzymatic and mechanical signaling cascade (e.g. CPC - RSs - I1 dynein/N-DRC – dyneins; Omoto and Kung, 1979; Smith and Yang, 2004; Brokaw, 2009), the geometric clutch model that suggests bending-induced distortions of the axoneme change the spacing between DMTs, acting as a “clutch” to disengage dyneins from their DMT tracks (Brokaw, 2009; Lindemann, 1994, 2007; Lindemann and Lesich, 2010), and other mechanical feedback models where beating causes deformations or stress (e.g. interdoublet sliding, normal forces or curvature of the flagellum) that in turn regulate motor activity (Sartori et al., 2016). Considering how many regulatory complexes have been identified within the 96-nm axonemal repeat, it is not unlikely that a combination of regulatory mechanisms controls ciliary motility.

Interestingly, our data revealed an even larger and functionally modulated connectivity between dyneins and major regulatory complexes than previously appreciated, which supports a distributor model (Omoto and Kung, 1979; Smith and Yang, 2004; Brokaw, 2009). The regulatory complexes I1 dynein, I1-tether, and N-DRC undergo substantial conformational changes (Figures 4, 5, and S3) that correlated with bend direction similar to the IDAs *a, g,* and *d*. We found state-specific attachments and dissociations between these regulators and dyneins; for example the I1 dynein stalks sometimes connected directly to the proximally located nexin linker, and the pre-powerstroke 1β-head connected through the I1 tether and tether ridge to RS3 and CSC (Figures 4A and 4C, Pre; Movie S5), which in turn are linked to the N-DRC, RS2 and some IDAs (Lin et al., 2012a). These connections provide potential paths for regulatory signal and/or feedback transduction. The regulatory importance of the 1β-HC motor domain was further highlighted by the fact that in mutants missing the 1β-HC motor domain IC138 was hyperphosphorylated (Figures 4G and 4H). This is an “off” state also observed in CPC/RS mutants (Bower et al., 2009; Hendrickson et al., 2004) that results in severe motility defects and the failure to regulate microtubule sliding *in vitro* (Toba et al., 2011).

Our observations of positional changes between DMTs in active flagella are also consistent with the geometric clutch model (Lindemann, 2007). We observed subtle, bend-dependent changes in the relative positions of neighboring DMTs predominantly in the regions of targeted inhibition (Figure 2D). Even though the positional changes were small, they might be sufficient to disengage dyneins from the neighboring DMT in a clutch-like manner, either by increasing interdoublet distance or by changing the angle between dynein stalk and microtubule interface so that the binding affinity of dynein for the DMT is decreased. The axoneme distortion on the inhibited side could be caused by slight bending (Sartori et al., 2016) and/or the unique position of the N-DRC_3 conformation that showed unique contacts to IDAs (Figure 5).

Built-in mechanical constraints certainly contribute to the formation of ciliary bending. For example, it is not trivial to generate (quasi)planar waveforms by applying tangential forces to the periphery of a cylinder (which should cause twisting of the cylinder). In sea urchin sperm flagella, the permanent bridge structures between the two CPC microtubules (Carbajal-Gonzalez et al., 2013) and DMT5/6 (Lin et al., 2012b) most likely prevent sliding between these microtubules, forming two rigid planes that mechanical constrain the preferred beating direction perpendicular to these planes. A contribution of mechanical feedback to trigger conformational switching at the end of a bend is also possible, but faces the “chicken-and-egg problem”, meaning if the mechanics of bends trigger switching, what initiates bending in the first place. In virtually all cilia and flagella bending initiates at their base and propagates to the tip. One possibility is that the previously shown structural and compositional specializations in the proximal region of flagella, such as minor dynein isoforms (Yagi et al., 2009), and/or asymmetric and doublet-specific distribution patterns of IDAs are built-in asymmetry generators between dynein forces on opposing side of the axoneme. This could lead to random bend initiation, similar to the random twitching of the paralyzed *pf19* axonemes, but at an inherently higher rate and with functional regulators that can then perpetuate the beating.

## AUTHOR CONTRIBUTIONS

D.N. conceived and directed the study. J.L. performed the experiments. J.L. and D.N. analyzed the data and wrote the manuscript.

## ACKNOWLEDGEMENTS

We thank Daniel T.N. Chen and Zvonimir Dogic (Brandeis University) for providing sea urchin sperm and suggestions on ATP reactivation of axonemes, Chen Xu for providing EM training and management of the electron microscopy facility at Brandeis University, Mary Porter (University of Minnesota) for providing the anti-IC138 antibody and for critically reading the manuscript, Khuloud Jaqaman (UT Southwestern Medical Center) for experimental suggestions, and the team from XVIVO Scientific Animations for Movie S6. We are also grateful to William J. Snell and Mike Henne (UT Southwestern Medical Center), Cynthia Barber, and Jerry H. Brown for critically reading the manuscript. We also thank Peter Satir, Ian Gibbons and others in the cilia field for their pioneering studies of ciliary motility. This work was supported by funding from the National Institutes of Health (GM083122 to DN) and March of Dimes Foundation (to DN). The authors declare no competing financial interests.

## STAR Methods

## CONTACT FOR REAGENT AND RESOURCE SHARING

Further information and requests for resources and reagents should be directed to and will be fulfilled by the Lead Contact, Daniela Nicastro.

## EXPERIMENTAL MODEL AND SUBJECT DETAILS

Adult male sea urchins (*Strongylocentrotus purpuratus*) were purchased from Monterey Abalone, Monterey, CA. *Chlamydomonas reinhardtii* strains used in this study include WT (wild-type strain: CC-125, 137c mt+), pWT (a pseudo-WT strain: *pf2-4::PF2-GFP*) (Heuser et al., 2009), a CPC-lacking mutant *pf19* (strain: cc-1037 mt+) (Witman et al., 1978), and two I1 dynein mutant strains: *pf9-3 (+α tail)* (strain: *pf9-3 G41a*) and *idα2-7 (+β tail)* (strain: *Idα2- 7::pCAP1*). The *pf9-3 (+α tail)* and *idα2-7 (+β tail)* strains lack the motor domains of 1α- and 1β-HC, respectively (Myster et al., 1999; Perrone et al., 2000). *Chlamydomonas* cells were grown in liquid Tris acetate/phosphate medium at room temperature with a light/dark cycle of 16:8 h.

## METHOD DETAILS

### Specimen preparation

Spawning of male adult sea urchins was induced by the injection of 1-2 ml of 0.5 M KCl into the perivisceral cavity (Gibbons, 1982). Sperm samples were collected and a small aliquot was diluted in artificial seawater (360 mM NaCl, 50 mM MgCl_2_, 10 mM CaCl_2_, 10 mM KCl, and 30 mM HEPES, pH 8.0) to examine the motility by light microscopy using the differential interference contrast (DIC) mode of a Marianas spinning disk confocal system (3I, Denver, CO) consisting of a Zeiss Axio Observer Z1 microscope (Carl Zeiss, Jena, Germany) equipped with a Yokogawa CSU-X1 spinning disk confocal head (Yokogawa, Tokyo, Japan) and a QuantEM 512SC EMCCD camera (Photometrics, Tucson, AZ). All harvested sperm cells were motile (Movie S1), and the samples were then divided to prepare three different types of samples: (part A) active flagella, (part B) ATPase-inhibited immotile flagella (Immotile-I), and (part C) demembranated immotile axonemes (Immotile-D). Part A was diluted in artificial seawater and rapidly frozen (as described below). Part B was diluted in artificial seawater containing the ATPase inhibitor erythro-9-(2-Hydroxy-3-nonyl)adenine hydrochloride (EHNA hydrochloride, 2 mM; Santa Cruz Biotechnology) (Bouchard et al., 1981); after incubation for five minutes, we confirmed by light microscopy that the sperm were completely immotile (Movie S2) and then rapidly frozen the sample. Part C was diluted in demembranation buffer (30 mM HEPES pH 8.0, 150 mM KCl, 4 mM MgCl_2,_ 0.5 mM EGTA, and 0.1% Triton X-100) to remove the flagellar membrane (but axonemes remained attached to the cell body). After incubation for one minute, the sperm were collected by centrifugation at 1000x g, resuspended in demembranation buffer (but without Triton X-100), and rapidly frozen.

Axonemes were isolated from *Chlamydomonas reinhardtii* cells as previously described (Heuser et al., 2012a). Briefly, flagella were detached from the cells using the pH-shock method (Witman et al., 1972) and purified by two centrifugation steps over 20% sucrose cushions. Purified flagella were demembranated with 1% IGEPAL^®^ CA-630 (Sigma-Aldrich, St. Louis, MO), and axonemes were collected by centrifugation at 10,000 x g for 10 min. Except for WT and *pf19* axonemes that were used for an ATPase assay, and pWT axonemes that were used for cryo-ET analysis, the axoneme pellet was directly dissolved in two-dimensional electrophoresis (2DE) lysis buffer (7 M urea, 2 M thiourea, 4% (wt/wt) CHAPS, 65 mM DTT, and 2% (vol/vol) IPG buffer (pH 3-10NL; GE Healthcare)) by vigorously stirring for 0.5 h. Cell debris and insoluble material were removed by centrifugation at 45,000 g for 1 h. The supernatant was aliquoted and stored at -70°C until subsequent analysis.

For the phosphorylation analysis of IC138, the protein samples dissolved in 2DE lysis buffer were precipitated using the 2-D Clean-Up Kit (GE Healthcare) and resuspended in Milli-Q water to approximately 4 mg/ml. Phosphatase treatment using Lambda Protein Phosphatase was performed as previously described (Yamagata et al., 2002; Lin et al., 2011) with minor modifications. Briefly, two 85-μl aliquots of the protein sample were mixed with 10 μl of 10% SDS and were vigorously vortexed for 20 s, followed by the addition of 695 μl of Milli-Q water, 100 μl of 10 mM MnCl_2_, 100 μl of 10× Lambda Protein Phosphatase buffer (New England Biolabs), and 10 μl of protease inhibitor cocktail (P9599, Sigma-Aldrich). To one of the two aliquots 1200 units of Lambda Protein Phosphatase (New England Biolabs) was added, and both samples aliquots were then incubated overnight at 30°C. Phosphatase-treated and untreated protein samples were precipitated using acetone (-20°C), resuspended in 2DE lysis buffer, further purified using the 2-D Clean-Up Kit (GE Healthcare) and resuspended in 2DE lysis buffer to a final concentration of approximately 4 mg/ml.

### Cryo-sample preparation and cryo-ET

Cryo-samples were prepared, imaged by cryo-ET and processed as previously described (Lin et al., 2014). Briefly, Quantifoil holey carbon grids (Quantifoil Micro Tools GmbH, Germany) were glow discharged, coated with 10-nm gold (Sigma-Aldrich), and loaded onto a homemade plunge-freezing device. 3 μl of sample (i.e. actively swimming sea urchin sperm cells, *Chlamydomonas* WT cells, immotile sea urchin sperm cells (demembranated or treated with EHNA), paralyzed *Chlamydomonas pf19* cells, isolated *Chlamydomonas* WT flagella or pWT axonemes, respectively) and 1 μl of five-fold concentrated BSA coated 10-nm colloidal gold solution (Iancu et al., 2006) were applied to the grid, blotted with filter paper for 1.5–2.5 seconds, and immediately plunge-frozen in liquid ethane. Vitrified specimens were transferred into a Tecnai F30 transmission electron microscope (FEI, Hillsboro, OR) with a cryo-holder (Gatan, Pleasanton, CA). Flagella/axonemes that appeared well preserved by EM inspection were imaged at 300 keV, with -6 μm or -8 μm defocus, under low-dose conditions and using an energy filter in zero-loss mode (Gatan, Pleasanton, CA) (20-eV slit width). Tilt series were recorded while stepwise rotating the sample from about -65 to +65° with 1.5–2.5° increments using the microscope control software SerialEM (Mastronarde, 2005). The cumulative electron dose per tilt series was limited to approximately 100 e/Å^2^. All images were digitally recorded on a 2k×2k charge-coupled device camera (Gatan, Pleasanton, CA) at a nominal magnification of 13,500, resulting in a pixel size of approximately 1 nm.

### Image processing

The tilt series images were reconstructed into 3D tomograms by weighted back projection using the IMOD software package (Kremer et al., 1996). Some tomograms were previously utilized for the analysis of axonemal dyneins (Lin et al., 2014). Only tomograms of intact and non-compressed flagella/axonemes were used for further data analysis. To enhance the signal-to-noise ratio and improve the resolution, subtomograms that contained the 96-nm axonemal repeat units along the doublet microtubules (volume size: 110 × 84 × 80 nm) or that contained individual ODAs (volume size: 56 × 56 × 56 nm) were extracted from the raw tomograms, aligned, and averaged (including missing wedge compensation) using the PEET program to obtain subtomogram averages (Nicastro et al., 2006). To identify distinct conformations of various axonemal structures, classification analyses was performed on the aligned subtomograms using a clustering approach (principle component analysis) built into the PEET program (Heumann et al., 2011). Prior to classification, appropriate masks were applied to focus the classification on structures of interest. Subtomograms that contained the structure of interest with an identical conformation were grouped into a class, and were averaged to generate a class average. The automatic classification into different conformational states of axonemal complexes was performed by an automated algorithm without prior knowledge about from which functional region of a flagellar wave individual subtomograms were extracted. Using the information provided by the classification analysis, we then mapped the conformational state of each subtomogram/repeat back onto the respective location in the raw tomograms. The numbers of tomograms and subtomograms analyzed by classification are summarized in Table S1. The resolution of the resulting averages was estimated in a (30 nm)^3^ subvolume in the center of the structure of interest using the Fourier shell correlation method with a criterion of 0.5 (Harauz and Van Heel, 1986). The structures were visualized as 2D tomographic slices and 3D isosurface renderings using IMOD (Kremer et al., 1996) and UCSF Chimera (Pettersen et al., 2004), respectively.

### Electrophoresis and phosphorylation analysis

2DE analysis was performed as previously described (Lin et al., 2011). Briefly, *Chlamydomonas* axonemal proteins (70 μg) were separated in the first dimension on 13-cm immobilized pH 3-10NL IPG strips (GE Healthcare) for 24 kV-h, followed by 10% SDS-PAGE for the second dimension. All samples (2-7 samples for each strain) were run in at least duplicate to confirm reproducibility. To visualize total proteins, the gels were stained with silver nitrate; to visualize phosphoproteins, the gels were stained with Pro-Q Diamond Phosphoprotein Gel Stain (Thermo Fisher Scientific) according to the manufacturer’s instructions. Following image acquisition using a Typhoon 9410 Variable Mode Imager (GE Healthcare), the Pro-Q Diamond-stained gels were post-stained with SYPRO^®^ Ruby Protein Gel Stain (Thermo Fisher Scientific) to detect total protein.

### Immunoblot

For 2D immunoblot analysis of IC138, 35 μg of total axonemal proteins was separated by 2DE with 7-cm immobilized pH 3-10NL IPG strips (GE Healthcare) for the first dimension, and 10% SDS-PAGE gels for the second dimension. Immunoblot analysis was performed using a polyclonal IC138 antibody (1:10,000) (Hendrickson et al., 2004). Signals were visualized using the ECL detection system (Bio-rad).

For immunoblot analysis of ODA IC2 protein, the axonemal proteins were resolved by SDS-PAGE on Any kD TGX Strain-FreeTM protein gels (Bio-rad Cat# 4568123). After visualizing by ChemiDocTM Touch Imaging System (Bio-Rad), proteins were blotted onto PVDF membranes (Bio-Rad) and probed with anti-IC2 monoclonal antibody (Sigma, D6168; 1:5000). Signals were visualized using the ECL detection system (Bio-rad), and quantified by ImageJ software.

### ATPase assay

The rate of the phosphate releases by ATP hydrolysis of axonemes was measured in bulk as described before (Chifflet et al., 1988). Briefly, isolated axonemes were washed and resuspended in HMEEK buffer (30 mM Hepes, pH 7.4, 5 mM MgSO_4_, 1 mM EGTA, 0.1 mM EDTA, and 25 mM KCl). 150 μl of axonemes (10.3 μg) were incubated with 1.3 μl of 50 mM ATP for 1 min. The ATP hydrolysis was stopped by adding 150 μl of 12% SDS solution. Color was developed by incubation with 300 μl of a 1:1 solution of 6% ascorbic acid in 1 N HCl and 1% (NH_4_)_6_Mo_7_O_24_.4H_2_O in 12% SDS for 10 min, and was stopped by adding 450 μl of 2% sodium citrate, 2% NaAsO_2_, and 2% acetic acid followed by 20 min incubation at room temperature. The concentrations of the released phosphate were calculated from the color absorbance at 850 nm. The assays were performed on three independent samples.

## QUANTIFICATION AND STATISTICAL ANALYSIS

The quantification and statistical analyses for cryo-ET data and subtomogram averages are integral parts of the software packages used. The local resolutions of class averages were determined by Fourier shell correlation (implemented in PEET software; 0.5 criterion). For the ATPase assays mean values and the standard deviation from three independent samples were calculated, and the data were analyzed by Student’s *t* test.

## DATA AND SOFTWARE AVAILABILITY

All data are available upon request. SerialEM is a software for automated EM data acquisition. IMOD and PEET are software packages for reconstruction of 3D tomograms and sub-tomogram averaging, respectively. ImageJ is widely used for analysis of Western blot data. All these software packages are free for academic laboratories. The 3D averaged structures of all major classes of the ODA, I1 dynein, and I1-tether have been deposited in the Electron Microscopy Data Bank (EMDB) under ID codes XXX (accession numbers in progress).

## Supplemental figure legends

**Table S1.**
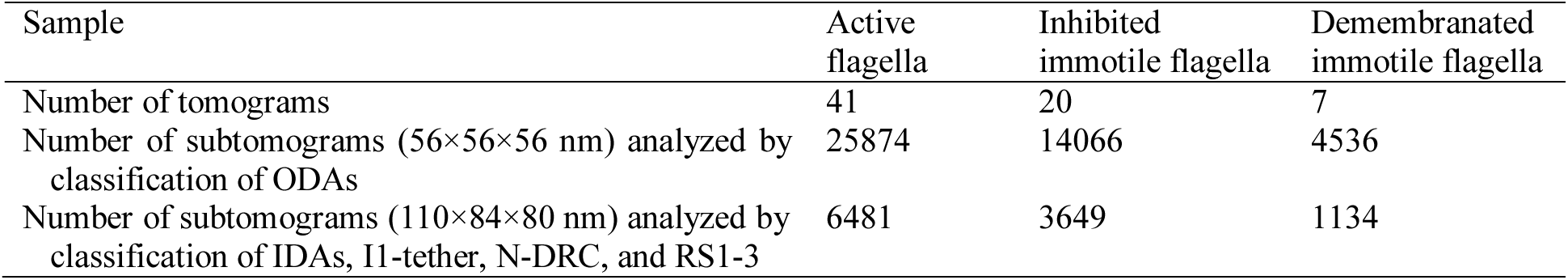
Summary of sea urchin sperm data analyzed by classification. Related to Figures 2-5.

| Sample | Active flagella | Inhibited immotile flagella | Demembranated immotile flagella |
| --- | --- | --- | --- |
| Number of tomograms | 41 | 20 | 7 |
| Number of subtomograms (56×56×56 nm) analyzed by classification of ODAs | 25874 | 14066 | 4536 |
| Number of subtomograms (110×84×80 nm) analyzed by classification of IDAs, I1-tether, N-DRC, and RS1-3 | 6481 | 3649 | 1134 |

**Table S2.**
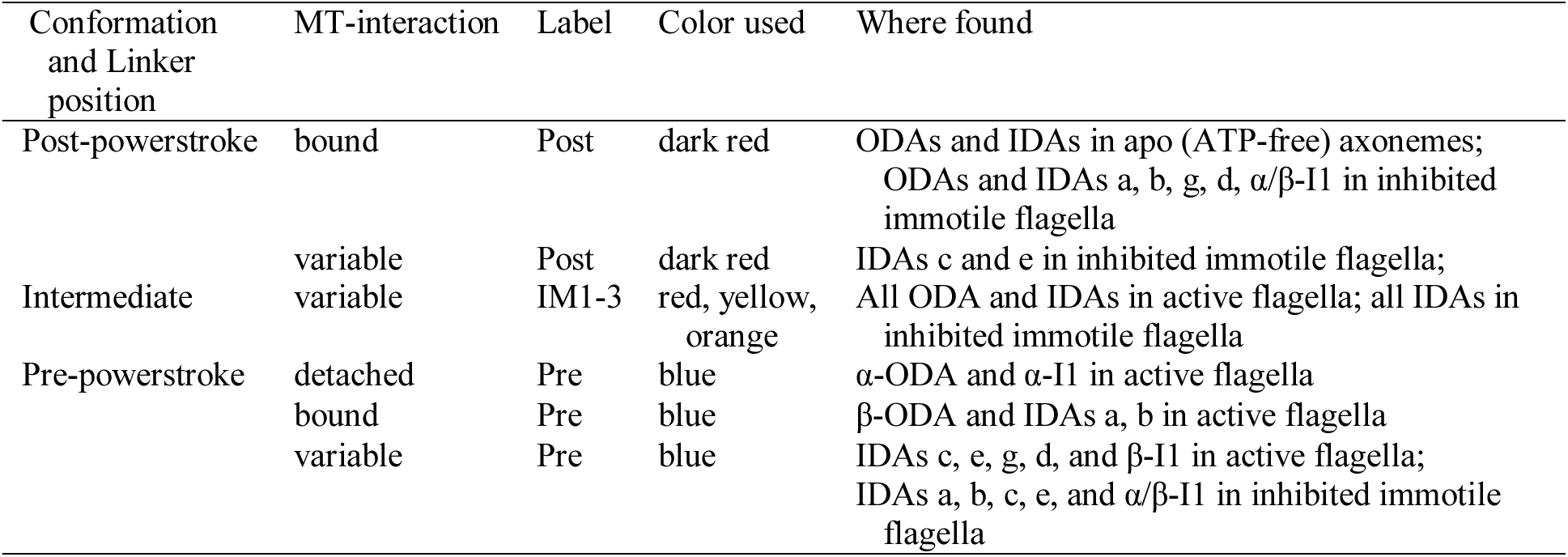
Summary of conformations of axonemal dyneins. Related to Figures 2-4.

**Figure S1.**
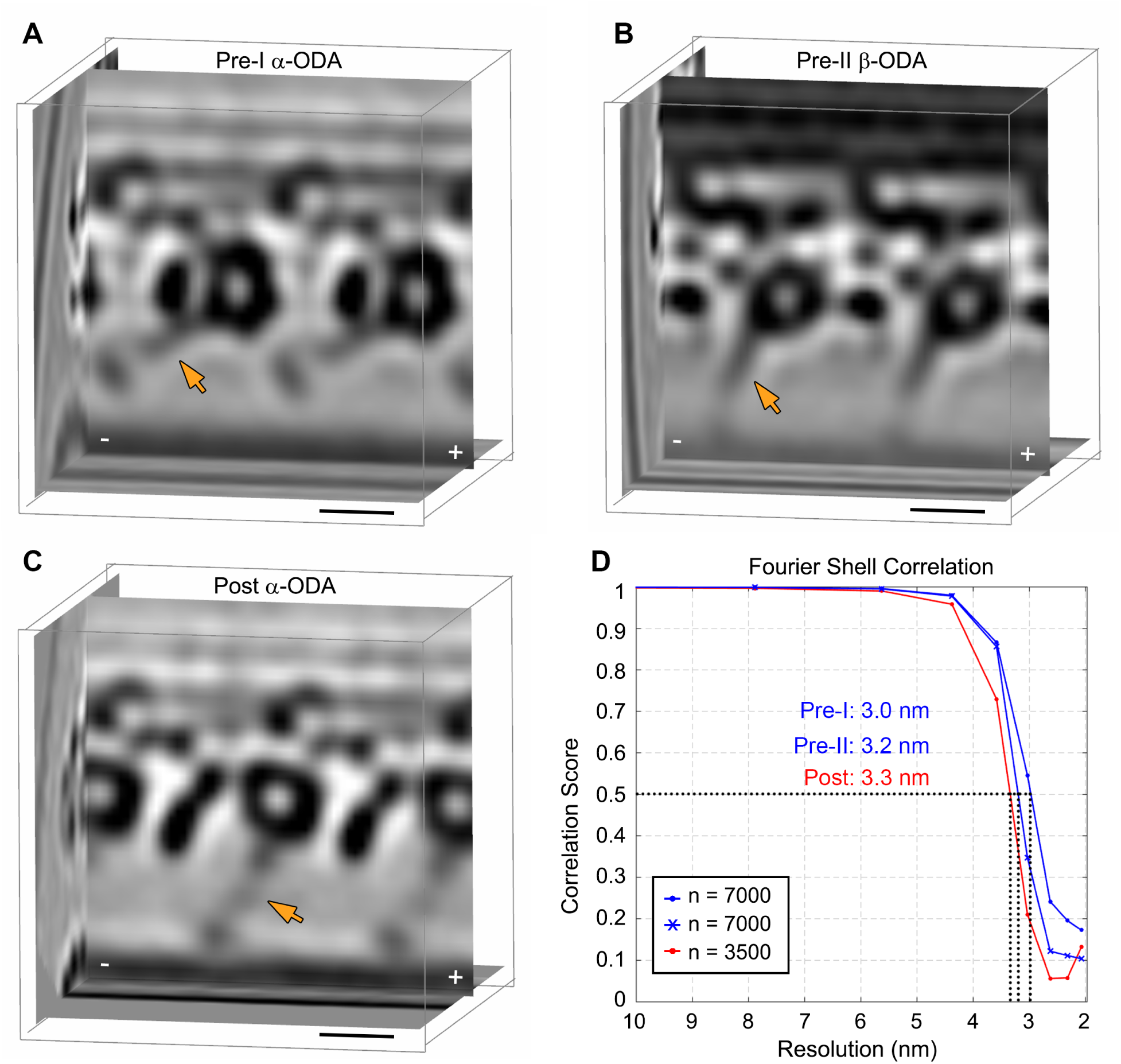
The predominant ODA conformations and their resolutions in active flagella and inhibited immotile flagella of sea urchin sperm. Related to Figure 2. (A-C) Tomographic slices of the class averages of outer dynein arms (ODAs) in the prepowerstroke Pre-I (A) and Pre-II (B) states in active flagella, and in the post-powerstroke conformation (Post) (C) in inhibited immotile flagella. The yellow arrows highlight the stalk of the ODA dyneins that is microtubule-detached in the primed Pre-I position (A), and microtubule-bound in both the primed Pre-II (B) and the Post conformation (C) (Lin et al., 2014). Microtubule polarity is indicated by -/+. (D) Spatial resolution of the three class average shown in (A-C) as determined by Fourier shell correlation (FSC=0.5). Scale bars: (A-C) 10 nm.

**Figure S2.**
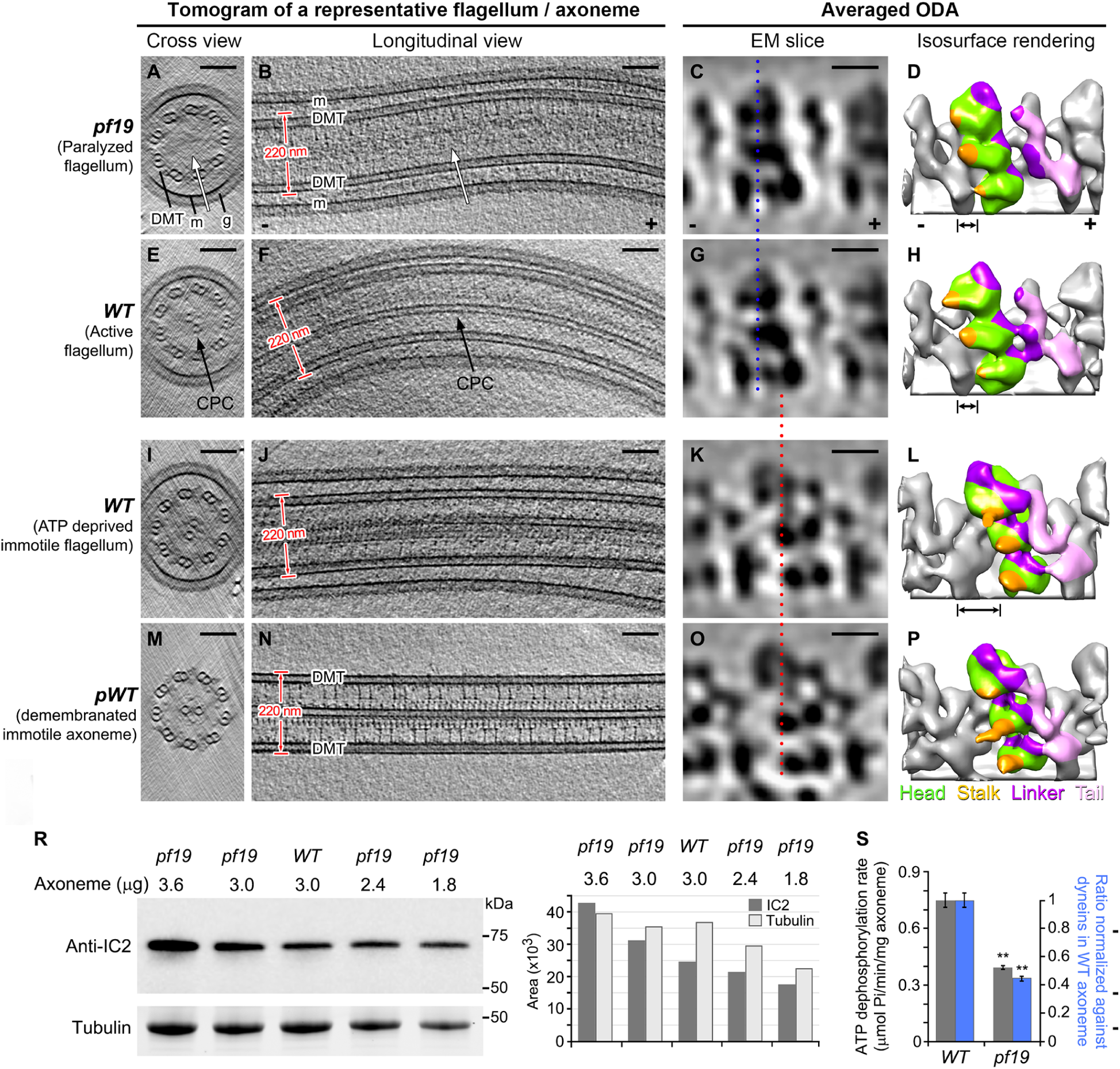
ODA dyneins in paralyzed *Chlamydomonas pf19* mutant flagella predominantly show pre-powerstroke conformation. Related to Figure 2. (A-P) Representative tomograms and averaged ODA conformations in the following *Chlamydomonas* samples: paralyzed flagella of intact *pf19* cells (A-D), active flagella from intact WT cells (E-H), detached WT flagella (I-L), and demembranated axonemes from pseudo-WT strain (*pf2-4::PF2-GFP*; biochemically, structurally and phenotypically indistinguishable from WT) (M-P). The latter two samples of isolated flagella/axonemes (I-P) were immotile, because detachment/demembranation allowed dilution/removal of ATP. The four columns show: 100-nm-thick cross-sectional (A, E, I and M) and 15-nm-thick longitudinal (B, F, J, and N) tomographic slices of a representative flagellum/axoneme; and 10-nm-thick longitudinal tomographic slices (C, G, K, and O) and 3D isosurface renderings (D, H, L, and P) of the averaged ODAs. Note the flagellar membrane (m) in (A, B, E, F, I and J) that is surrounded by a dense glycocalyx (g) layer that resulted in thicker samples and noisier tomograms for the *Chlamydomonas* flagella samples compared to the demembranated axonemes (M and N). Note also the missing central pair complex (CPC) in *pf19* (white arrows in A and B). The averaged ODAs in both paralyzed *pf19* and active WT flagella were predominantly in their prepowerstroke conformations with the three dynein heads (green) and stalks (orange) closer to the minus (-) end of the doublet microtubule (DMT) (C, D, G and H) than in the post-powerstroke conformation observed in detached and demembranated flagella (K, L, O and P). The blue/red dotted lines along the proximal edge of bottom dynein head (in C, G, K, and O) and the double-ended arrows (in D, H, and L) facilitate comparison of the head positions. (R) To properly normalize the ATP consumption measurements shown in (S) to the number of dyneins, the total axonemal protein amount was not suited, because the *pf19* axonemes lack the large (dynein-free) central pair complex (CPC) compared to WT axonemes. Therefore we compared the relative abundance of the outer dynein light chain protein IC2 in WT and *pf19* axoneme samples by immunoblot analysis. Various amounts of axonemal proteins were separated on a Stain-free SDS-TGX-gel (left bottom), blotted to a PVDF membrane, and then probed with anti-IC2 antibody (left top). The densitometry quantification of the bands was performed by ImageJ software (right). Based on analyses of three independent samples, the abundance of IC2 compared to total protein was 1.19±0.03 times higher in *pf19* than in WT axonemes. (S) ATPase assays. The ATPase activities of the WT and *pf19* axonemes were measured in the presence of 0.43 mM ATP. The bar chart shows the data as means ± SD of three independent samples (**, *p* < 0.01; Student’s *t* test). For the same amount of axonemes, the ATPase activity of *pf19* is 0.53±0.02 of that of WT (gray bars). As the abundance of dyneins in *pf19* axonemes is 1.19 times of that in WT axonemes (see R), the ATPase activity of *pf19* is 0.44±0.02 of that of WT after accounting for the amount of dyneins and normalizing against WT (blue bars). Scale bars: (A, B, E, F, I, J, M, and N) 100 nm, (C, G, K, and O) 10 nm.

**Figure S3.**
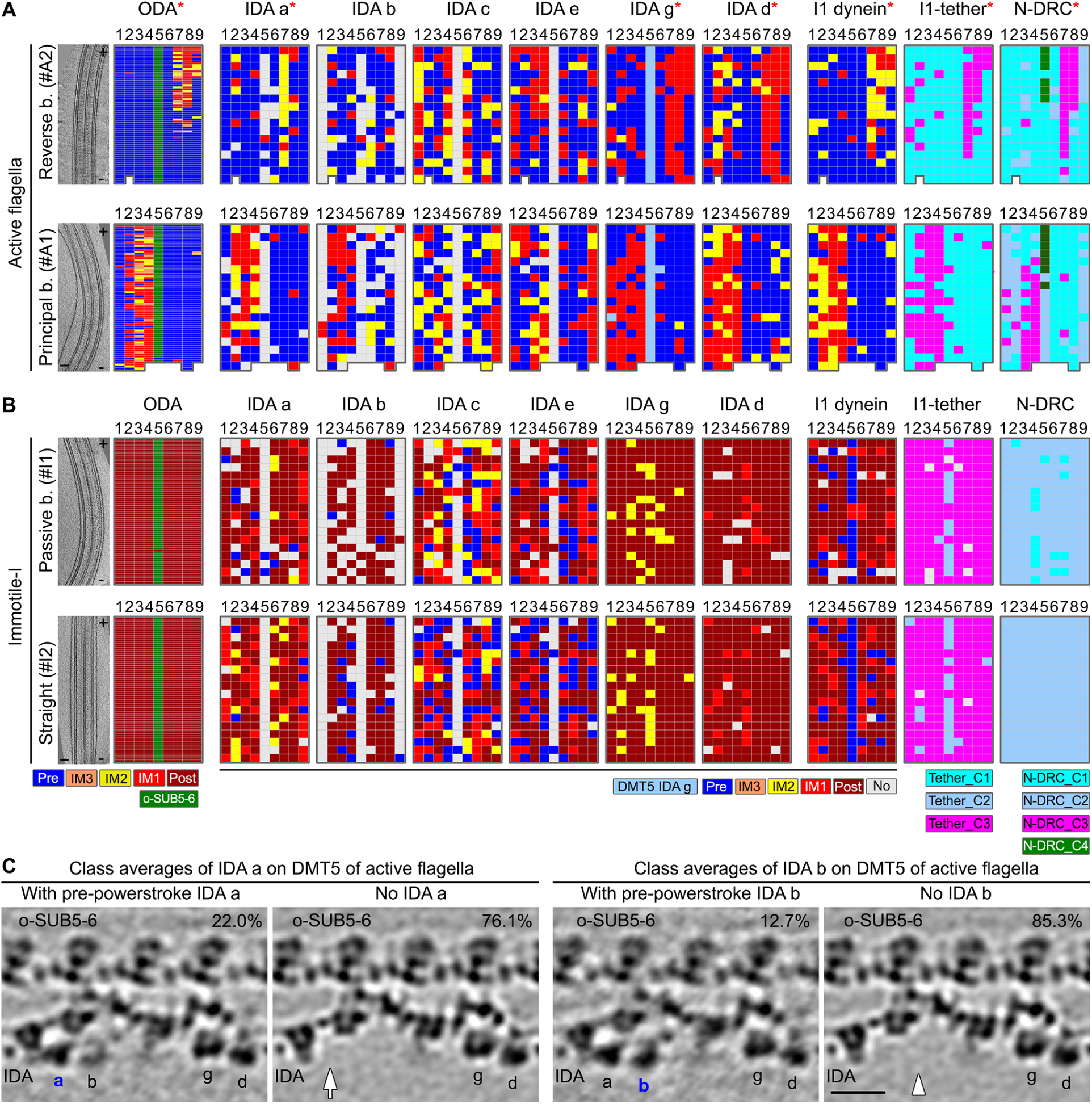
Classification analysis of dyneins, I1-tether, and N-DRC. Related to Figures 2-6. (A and B) For individual active flagella (A) and inhibited immotile flagella (B), a longitudinal tomographic slice (left) and the distribution patterns of conformations for various axonemal structures are shown. The structures on the nine DMTs (1-9; microtubule polarity indicated by +/-) are schematically shown as individual grids; the grid color represents the conformational state of each structure (as indicated in the bottom legends). Note that in immotile inhibited flagella (Immotile-I) (B), a small number of inner dynein arms (IDAs) showed pre-powerstroke or intermediate conformations, which might be due to incomplete inhibition by EHNA. However, unlike in active flagella (A), these Pre and IM conformations did not show a bend-correlated distribution pattern. (C) Longitudinal tomographic slices of class averages of IDA a (left) and IDA b (right) on DMT5. The arrow and arrowhead indicate the absence of IDAs a and b, respectively. The percentage of sub-tomograms included in each class average is indicated on the top right of each panel. Note that the absence of IDA a and b from most 96-nm repeats on DMT5 agrees well our previous study using doublet-specific averaging (Lin et al., 2012b); the successful detection of a small numbers of 96-nm repeats on DMT5 that did contain IDA a and/or b validates the reliability of the automatic classification method. Scale bars: (A and B) 100 nm, (C) 20 nm.

**Figure S4.**
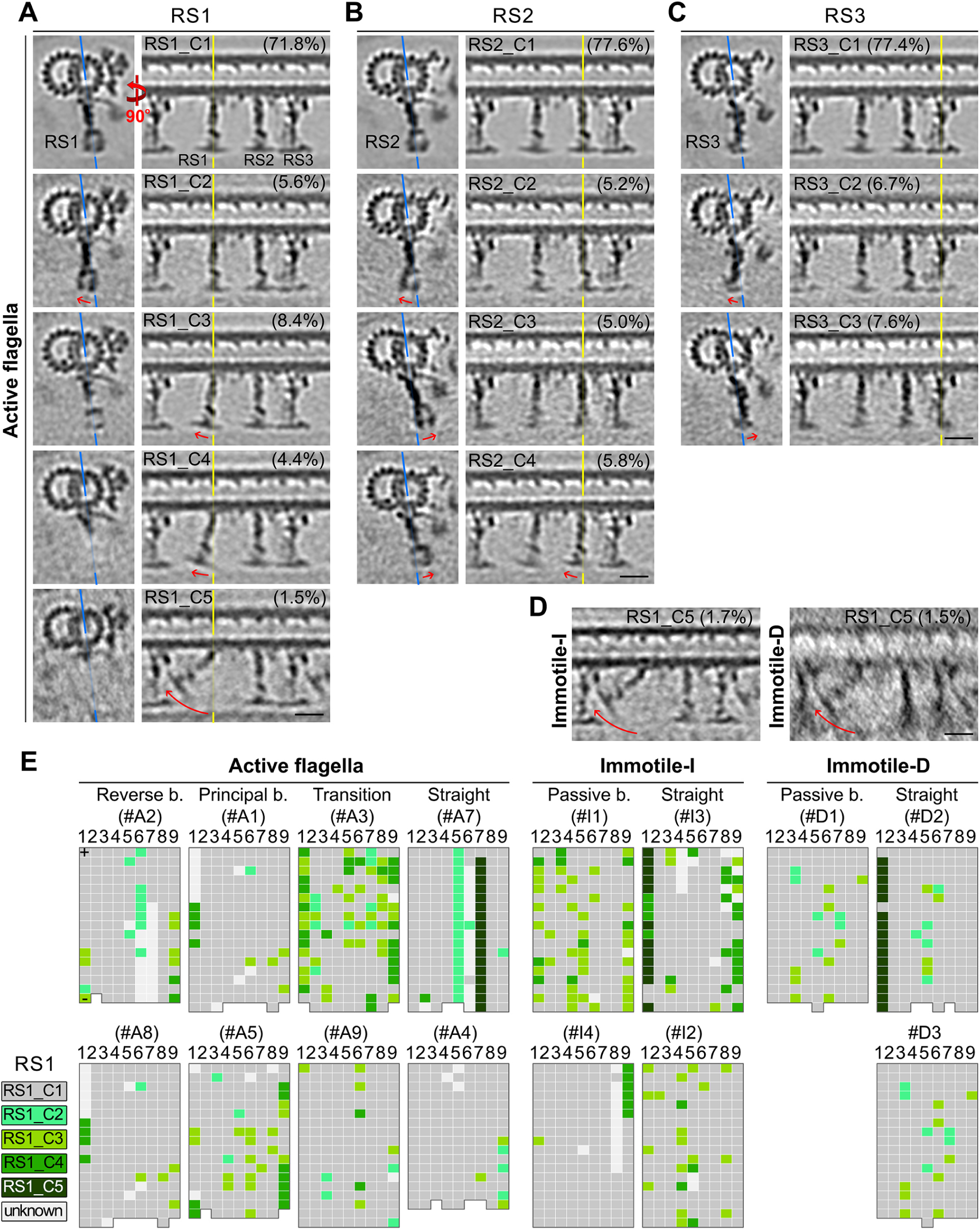
Tilted RSs are observed in both bent and straight regions of active flagella, as well as in immotile control samples. Related to Figures 4 and 6. (A-C) Tomographic slices of class averages of radial spokes RS1 (A), RS2 (B), and RS3 (C) in active flagella in cross-sectional (left) and longitudinal (right) views. Blue and yellow lines indicate the locations of longitudinal and cross-sectional slices, respectively. The percentage of sub-tomograms included in each class average is indicated on the top right. Red arrows highlight the tilts relative to class 1 (“_C1”) of each RS. Note that we observed the largest range of tilts for RS1; a possible explanation for this is that the spoke head of RS1 (horizontal density at the bottom of the spokes) in sea urchin sperm (and mammalian) flagella does not interact with neighboring spokes, whereas the spoke heads of RS2 and RS3 interact with each other (Lin et al., 2012a), limiting their range of tilting. (D) Tomographic slices of class averages of RS1_C5 in inhibited immotile flagella (Immotile-I) and demembranated immotile axonemes (Immotile-D) show the same large tilt of RS1 that was also observed for RS1_C5 in active flagella. The percentage of sub-tomograms included each class average represents is indicated on the top right. (E) Distributions of RS1 class averages with different degrees of tilt in different regions of individual active, immotile-I, and immotile-D flagella. RS1s on the nine DMTs (1-9) are schematically shown as individual grids; the grid color represents the tilt state of each RS1 (as indicated in the legend on the left). Tilting of radial spokes was previously indicated as possibly regulatory mechanism (Warner and Satir, 1974). However, we observed RS tilting in both active and immotile flagella without clear correlate to the bending direction of functional regions of the sinusoidal wave. Scale bars: (A-D) 20 nm.

## Supplemental movie legends

**Movie S1. DIC light microscopy movie of swimming sea urchin sperm cells.** Related to Figure 1.

The actively beating flagella of sea urchin sperm that were freshly harvested and diluted in artificial seawater, drive the rapid swimming motion of the sperm cells. This movie is reproduced from (Lin et al., 2014).

**Movie S2. DIC light microscopy movie of inhibited and thus immotile sea urchin sperm cells.** Related to Figure 1.

The motility of sea urchin sperm flagella is completely inhibited by diluting the freshly harvested sperm cells in artificial seawater containing the (dynein) ATPase inhibitor EHNA (Bouchard et al., 1981). Only passive drifting movement of the sperm cells due to buffer flow was observed. This movie is reproduced from (Lin et al., 2014).

**Movie S3. Using cryo-ET, subtomogram averaging and classification analysis of active sea urchin sperm flagella, we identified distinct ODA conformations that show a bend-correlated distributions inside the flagella.** Related to Figures 2 and 6.

The classified ODA conformations were mapped to their location in the raw tomograms of the bent flagellum. The animation starts with sequential tomographic slices through the principal bend region of an active flagellum (oriented with its proximal side to the right); this is followed by an overlaid, simplified graphical model in which small colored spheres show the locations of ODA in the following conformations: pre-powerstroke (*Pre*, blue), intermediate states (*IM3*, orange, *IM2*, yellow, and *IM1*, red), and o-SUB5-6 bridge (green). The intermediate conformations (IM1-3) are mainly present on DMTs 3 and 4. For a principle bend the dyneins on these DMT are predicted to be inactive according to the previous “switch-point” hypothesis (Satir and Sale, 1977; Wais-Steider and Satir, 1979).

**Movie S4. IDAs c and e exhibit large conformational changes in active flagella.** Related to Figures 3 and 6.

The animation switches between four longitudinal tomographic slices of class averages of IDAs c and e, showing the conformational changes that these two IDAs undergo during flagellar beating. The general location of the tomographic slices is the same as in Figure 3B. The four class averages are shown in the following sequence: IM1 (IDAs c and e); IM2 (IDAs c and e); Pre (IDAs c and e); and Pre (IDA c) and IM1 (IDA e) (see also Figure 3). Note that these two dynein isoforms did adopt different conformations in the active flagella, but showed no obvious bend-correlated distribution pattern; the sequence of depicted conformational changes was chosen randomly and does not imply a particular sequence of states. Scale bar: 20 nm.

**Movie S5. The two motor domains of the I1 dynein and the associated tether and tetherhead exhibit large conformational changes in active flagella.** Related to Figures 4 and 6.

The animation switches between the isosurface renderings of the pre-powerstroke (Pre) and the intermediate class IM2 averages of I1 dynein, showing a rotation of the I1 dynein heads around each other. The animation starts with the IM2 conformation, in which 1β-motor domain is detached from the ridge density of the I1-tether, followed by the Pre conformation, in which 1β-motor domain is attached to the ridge density. Correlated to the I1 dynein conformational changes, the I1-tether also shows large conformational changes.

**Movie S6. The animation summarizes the switch-inhibition mechanism of ciliary and flagellar beating, as suggested by our cryo electron tomography study of sea urchin sperm flagella.** Related to Figures 2-6.

KEY RESOURCES TABLE

KEY RESOURCES TABLE
| REAGENT or RESOURCE | SOURCE | IDENTIFIER |
| --- | --- | --- |
| **Antibodies** |  |  |
| Rabbit polyclonal anti-IC138 | Laboratory of Mary porter; Hendrickson et al., 2004 | N/A |
| Mouse monoclonal anti-IC2 | Sigma-Aldrich | Cat#D6168 |
| Goat Anti-Mouse IgG (H + L)-HRP Conjugate | Bio-rad | Cat#1706516 |
| Goat Anti-Rabbit IgG (H + L)-HRP Conjugate | Bio-rad | Cat#1706515 |
| **Chemicals, Peptides, and Recombinant Proteins** |  |  |
| erythro-9-(2-Hydroxy-3-nonyl)adenine hydrochloride | Santa Cruz Biotechnology | Cat#sc-201184A |
| TGX Strain-Free<sup>TM</sup> protein gels | Bio-rad | Cat#4568123 |
| Clarity Western ECL Substrate | Bio-rad | Cat#170-5060 |
| Hutner's Trace Elements | *Chlamydomonas* Resource Center | N/A |
| IGEPAL® CA-630 | Sigma-Aldrich | Cat#I8896-50ML |
| Protease inhibitor cocktail | Sigma-Aldrich | Cat#P9599 |
| ATP | Sigma-Aldrich | Cat#A2383 |
| 10-nm gold Gold nanoparticles | Sigma-Aldrich | Cat#752584 |
| 13 cm IPG buffer (pH 3-10NL) | GE Healthcare | Cat#17-6000-88 |
| Immobiline DryStrip (pH 3-10NL, 7 cm) | GE Healthcare | Cat#17-6001-12 |
| Immobiline DryStrip (pH 3-10NL, 13 cm) | GE Healthcare | Cat#17-6001-15 |
| 2-D Clean-Up Kit | GE Healthcare | Cat#80-6484-51 |
| Lambda Protein Phosphatase | New England Biolabs | Cat#P0753S |
| Pro-Q Diamond Phosphoprotein Gel Stain | Thermo Fisher Scientific | Cat#P33300 |
| SYPRO® Ruby Protein Gel Stain | Thermo Fisher Scientific | Cat#S12000 |
| **Deposited Data** |  |  |
| Sub-tomogram average of ODA Pre class | This study | (accession numbers in progress) |
| Sub-tomogram average of ODA Post class | This study | (accession numbers in progress) |
| Sub-tomogram average of I1 dynein Pre class | This study | (accession numbers in progress) |
| Sub-tomogram average of I1 dynein Post class | This study | (accession numbers in progress) |
| Sub-tomogram average of I1-tether Tether_C1 class | This study | (accession numbers in progress) |
| Sub-tomogram average of I1-tether Tether_C3 | This study | (accession numbers in progress) |
| **Experimental Models: Organisms/Strains** |  |  |
| Sea urchins: *Strongylocentrotus purpuratus* | Monterey Abalone, Monterey, CA | N/A |
| *Chlamydomonas* WT (strain: CC-125, 137c mt+) | Laboratory of Mary porter | N/A |
| *Chlamydomonas* pWT(a pseudo-WT strain: *pf2-4::PF2-GFP*) | Laboratory of Mary porter; Heuser et al., 2009 | N/A |
| *Chlamydomonas pf9-3 (+$\alpha$ tail)* (strain: *pf9-3 G41a*) | Laboratory of Mary porter; Myster et al., 1999 | N/A |
| *Chlamydomonas ida2-7 (+$\beta$ tail)* (strain: *Ida2-7::pCAP1*) | Laboratory of Mary porter; Perrone et al., 2000 | N/A |
| *Chlamydomonas* pf19 (strain: CC-1037, mt+) | *Chlamydomonas* Resource Center | http://www.chlamycollection.org/product/cc-1037-pf19-mt/ |
| Software and Algorithms |  |  |
| SerialEM | Mastronarde, 2005 | http://bio3d.colorado.edu/SerialEM/ |
| IMOD | Kremer et al., 1996 | http://bio3d.colorado.edu/imod/ |
| PEET | Nicastro et al., 2006 | http://bio3d.colorado.edu/PEET/ |
| UCSF Chimera | Pettersen et al., 2004 | http://www.cgl.ucsf.edu/chimera/ |
| ImageJ | NIH | https://imagej.nih.gov/ij/ |
| Other |  |  |
| Holy carbon EM grid (Cu, R2/2, 200 mesh) | Quantifoil | http://www.quantifoil.com/index.php?name=circular |

## Footnotes

* Cilia and flagella share the same conserved structure and differ only in waveform and/or length; therefore the terms “cilia” and “flagella” will be used interchangeably in this paper.

